# WNK1 kinase activity is required for maintenance of podocyte foot process structure

**DOI:** 10.1101/2025.05.30.657136

**Authors:** Zhenan Liu, Eunyoung Lee, Shumeng Jiang, Joonho Yoon, Fouzia Ahmed, Mohammad A. Rahman, W. Philip Bleicher, Hani Y. Suleiman, Leslie A. Bruggeman, R. Tyler Miller, Audrey N. Chang

## Abstract

The filtration-function of glomeruli requires slit diaphragms formed by interdigitating podocyte foot processes, which are actin-based membrane protrusions. Dysregulation of mechanisms that maintain these membrane extensions lead to foot process effacement, proteinuria, and progression to chronic kidney disease. Building on our previous work that showed WNK1 kinase activity is necessary for the maintenance of normal biomechanical properties of glomeruli and podocyte foot process architecture, we tested the hypothesis that WNK1 kinase activity affects the structure of podocyte foot processes through modulation of actomyosin activity and focal adhesion complexes. Using a WNK1 kinase specific inhibitor, we determined by immunofluorescence microscopy of nascent focal adhesions, podocyte membrane spreading/extensions, and NMII paralog localization and extent of activation calculated from quantification of phosphorylated myosin, that all were sensitive to WNK1 kinase activity. Moreover, biochemical evidence of WNK1 kinase activity-dependent signalosomes supports a role for WNK1 in the maintenance of podocyte foot processes, and sarcomere-like structures (SLSs) that are induced in models of podocyte injury. Using primary and immortalized podocyte cell lines developed from control and *Col4a3^-/-^*Alport Syndrome model mice, we measured WNK1 kinase activity-dependent improvement in properties of injured podocytes *in vitro*. Physiological relevance of WNK1 kinase activity-dependent structural maintenance of podocyte foot processes was confirmed by significant acute proteinuria measured in response to WNK1 inhibition *in vivo*. Collectively, the results provide evidence that WNK1 kinase signalosome activity that includes formation of nascent focal adhesions and regulation of NMII localization and activity at membrane protrusions and extensions, are necessary for physiological maintenance of slit diaphragms.

**Significance:** Terminally differentiated podocytes are arborized cells with interdigitating foot processes that form the renal filtration barrier. Loss of foot process structural integrity causes progressive proteinuria, which can lead to irreversible renal injury, but the mechanisms that maintain foot process structure are incompletely understood. We report evidence that WNK1 kinase activity is required for maintenance of normal glomerular filtration *in vivo*, and this is mediated in part through WNK1 activity-dependent modulation of non-muscle myosin II activity, and formation of nascent focal adhesions that are necessary for lamellipodial extensions. Using glomeruli and podocyte cell lines developed from an Alport Syndrome podocyte injury model, we show that aspects of abnormal podocyte structure associated with chronic kidney disease can be suppressed through increased WNK1 activation.

## Introduction

Disrupted podocyte actin cytoskeletal structure due to mutations in structural or regulatory proteins or their dysfunction is the foundation of many glomerular diseases and disease models. Various *in vitro* models of podocyte injury demonstrate reduced capacity to spread and generate traction force[1], indicating baseline dysfunction in focal adhesion formation and/or actomyosin regulation. Extensive interdigitation of cell processes among neighboring podocytes forms the slit diaphragm, essential components of the cellular sieve that retain proteins in the blood while allowing filtrate to enter the urine for reabsorption or excretion[2]. A characteristic of kidney disease progression is loss of podocyte structural complexity and adhesion strength with resultant proteinuria, filtration failure, and podocyte loss[3,4].

Podocytes hypertrophy and remodel in response to neighboring podocyte loss or glomerular growth[5,6]. The maintenance of arborized podocyte structures requires coordinated formation and activation of focal adhesions for dynamic regulation of the cell cytoskeleton that are responsive to hemodynamic forces and matrix stiffness[7,8]. In order to retain glomerular filtration function while under pulsatile hemodynamic forces in a 3D environment, constant dynamic regulation of myosin activity at the membrane edges and in cell extensions is necessary for the formation and maintenance of the slit diaphragm, and in focal adhesions for adhesion to the basement membrane. In the 2D environment, podocytes in culture migrate as they grow and divide, relying on assembly of nascent focal adhesions at leading edges of lamellipodial extensions, which undergo dynamic maturation to focal complexes and focal adhesions[9].

While the structures of actin filaments and focal adhesions at lamellipodia extensions and stress fibers of the cell are well documented in various cell types including podocytes[10], mechanisms that underlie regulation of membrane dynamics have not been defined clearly. Extending our previous measurements of WNK1 kinase activity in podocyte membrane blebs[11], we sought to determine whether WNK1 activity-dependent regulation of membrane dynamics is mediated through WNK1-associated focal adhesion components that contribute to actomyosin regulation at lamellipodial extensions.

Non-muscle myosin IIs (NMIIs) are the contractile apparatus of actomyosin cytoskeletal complex where they hydrolyze ATP to convert chemical energy into mechanical functions in cells. Non-muscle myosin II has three paralogs named as NMIIA, NMIIB and NMIIC. They are essential proteins for sensation of and responses to mechanical force, control of cell shape, cell migration, and biomechanical properties. Relatively little work has focused on NMIIs in podocytes despite their importance in cell structure, possibly because the phenotypes of mutations in them and knockout mice are relatively subtle compared to those of other proteins. This apparent phenotypic subtlety may be a consequence of functional redundancy as well as the lack of paralog-specific inhibitors that can be used systemically to assess their role in organ or tissue function. Experimental manipulation of the NMIIs in isolated glomeruli and cultured cell systems demonstrates their importance in podocytes on a cell biologic level and supports physiological roles in glomerular disease.

NMIIs are ubiquitously expressed conventional type II myosins[12], the superfamily which includes distinct muscle type-specific myosins[13]. Conventional myosins are hexameric ATPases formed by dimerization of two protomers comprised of a heavy chain that has a motor domain “head” and elongated coiled-coil “tail”, and two light chains wrapped around the “neck” or lever-arm of the myosin molecule. The two light chain molecules are distinct gene products; the alkali or essential light chain (ELC) stabilizes the myosin neck, and the regulatory light chain (RLC) determines the orientation of the myosin head through a phosphorylation-dependent mechanism. NMIIs form bipolar bundles through tail-tail assembly, and bi-directional myosin ATPase activity drives motility. Cell polarity is driven in part by the differential distribution of NMII paralogs[14,15]. Proteomics (https://kidneyapp.shinyapps.io/podident/) and scRNAseq (humphreyslab.com) data show that podocytes express three NMII paralogs, *MYH9* (NMIIA), *MYH10* (NMIIB) and *MYH14* (NMIIC). The molar ratios of NMII paralogs are not reported for podocytes, but NMIIA is the best studied because it is the most abundant transcript and mutations in it are associated with glomerular disease such as focal segmental glomerulosclerosis (FSGS)[16,17]. In healthy podocytes, NMIIA was shown to be expressed mainly in the cytoplasm of primary processes of the arborized cell structure. In injured podocytes, NMIIA forms patterned sarcomere-like structures (SLSs) with actin and synaptopodin, juxtaposed to the glomerular basement membrane [17,18]. In contrast to NMIIA, NMIIB has not been extensively studied in the podocytes. Various reports that support a role for all NMII paralogs in cells, regardless of amount of protein, raise the importance of understanding the regulatory mechanisms that control NMIIs and underlie maintenance of podocyte structure[19,20].

NMIIs require phosphorylation of the RLC subunit for actin-activated ATPase reactions. Non- and smooth muscle RLC (Myl12a/b, Myl9) are mono- or di-phosphorylated at Thr18/Ser19 in human and Thr19/Ser20 in mouse. The canonical regulators of NMII ATPase reactions that drive contractions in non-muscle cells are the Ca^2+^/CaM-dependent myosin light chain kinase (MLCK) and the myosin light chain phosphatase (MLCP). The gene for MLCK in non-muscle cells, *MYLK*, encodes 3 distinct proteins: long non-muscle MLCK, short smooth muscle MLCK, and telokin[21]. Differential expression of long non-muscle and short smooth muscle MLCK in tonic and phasic smooth muscle tissues and cell types suggests distinct contributions of the two variants toward regulation of myosins[22]. Non-muscle MLCK is dominantly expressed in the kidney, consistent with observed enrichment in other non-muscle cells [23]. The MLCP that regulates inactivation of NMIIs is a holoenzyme comprised of the regulatory myosin target subunit MYPT1, a catalytic subunit PP1c, and an accessory protein M21[24]. MYPT1 is ubiquitously expressed and targets the catalytic subunit PP1c to myosin through high affinity binding sequences, to dephosphorylate RLC. Expression levels of these regulatory proteins, basal activation states, and localization within normal and diseased glomerular podocytes have not been clearly documented. Given the importance of NMIIs in determining cell structure and behavior, information on basic properties of actomyosin regulation in podocytes as it relates to focal adhesion formation, is necessary to understand its dysregulation in the context of podocyte injury.

With-no-lysine (WNK)1 kinase is expressed in glomerular podocytes and contributes to regulation of glomerular structure[11]. Chemical inhibition of WNK kinases reduced mouse glomerular stiffness, decreased F-actin, and disrupted podocyte foot process structure, causing a reduction in slit diaphragm density[11]. Conversely, activation of WNK1 kinase by calcineurin inhibitors FK506 or cyclosporin increased glomerular stiffness and F/G actin ratio. In cultured podocytes, WNK1 kinase activation increases podocyte membrane blebbing, lamellipodia formation, and cell migration and contractility[11], processes that all require NMII activity. These results point to a novel role for WNK1, a well-known regulator of ion transport in renal tubules[25] and the vascular system[26], in podocyte cytoskeletal structure. Herein, we investigated the distribution of NMII paralogs in mouse glomerular podocyte foot processes, and the membrane edges and lamellipodial extensions of podocytes in culture, to test the hypothesis that WNK1 kinase activity mediated NMII actomyosin activation at the leading edges of membrane extensions is required for maintenance of podocyte foot processes, and enhance our understanding of NMII paralog distribution as it relates to cell morphology and function.

Using isolated primary podocytes from glomerular outgrowths, we identified differential distribution of NMIIA and NMIIB at membrane blebs and lamellipodial extensions, and evidence of WNK1-activity dependent formation of WNK1 and vinculin association that contribute to NMII activation. Sarcomere-like patterning of synaptopodin and NMIIs in VRAD-differentiated WT podocytes, were disrupted with WNK1 inhibition, indicating its activity may be necessary for maintenance of not only membrane targeted NMIIs but also stress-induced cytoplasmic NMII adhesion mechanisms in pathological states. Moreover, cell lines developed from the *Col4a3^-/-^* model of Alport Syndrome, showed evidence of WNK1 signaling dysregulation, further supporting a role for WNK1 in renal pathology associated with podocyte injury. Acute proteinuria in response to WNK1 inhibition *in vivo*, supports a physiological role for WNK1 in maintenance of podocyte slit diaphragms. Collectively, these results provide evidence that WNK1 contributes to mechanosensation in podocytes that through formation of activity-dependent signalosomes, modulates NMII activity at nascent focal adhesions and at sarcomere-like structures in pathological states. Additionally, rescue of *Col4a3^-/-^* glomerular structure with acute WNK1 activation supports WNK1 activity regulation as a potential mechanism of action for calcineurin inhibitors’ beneficial effects in glomerular disease, and suggest further study of the effects of WNK1 kinase activity on podocytes and glomerular capillaries may lead to identification more specific targets.

## Results

### Differential expression of NMIIA and NMIIB in podocyte foot processes and lamellipodial extensions, where the WNK1 kinase substrates OSR1/SPAK are found

As previously reviewed[10], podocyte NMIIA has been studied in depth due to greater mRNA and protein expression levels relative to NMIIB, and its association with hereditary glomerular disease. While NMIIA is the main paralog expressed in the podocytes and is important for podocyte cell adhesion upon injury [18], less is known about the role of NMIIB [17,18,27]. NMIIB was reported to be undetectable in glomeruli, but is expressed in cultured podocytes with non-overlapping functions with NMIIA, where it contributes to cell adhesion, directly affecting cell spreading, the number of stress fibers, focal adhesion density, and cell attachment [27]. In a prior study, using immunofluorescence microscopy, we found that WNK1 and its substrate OSR-1 localized to synaptopodin-positive regions of mouse glomeruli [11]. As such, we asked whether WNK1 colocalized with NMIIA, and whether NMIIB was indeed excluded from synaptopodin-positive regions of the glomerulus.

Using antibodies that were validated with knockout tissues [19,28], we localized NMIIA and NMIIB in the glomerulus by immunofluorescence microscopy. Although the NMIIA and NMIIB could not be co-imaged at the same time because the antibodies were from the same species, comparisons of their expression within mouse glomeruli co-stained for synaptopodin showed NMIIA and NMIIB are both expressed in synaptopodin-positive regions of the isolated glomerulus (Figure 1A, arrows). Qualitative comparisons of Z-stack images (not shown) and super resolution microscopy of cryosectioned cortical tissue (Figure 1B) showed consistently greater co-localization of NMIIB and the podocyte-specific marker synaptopodin (yellow color in merged panel), than NMIIA.

**Figure 1.**
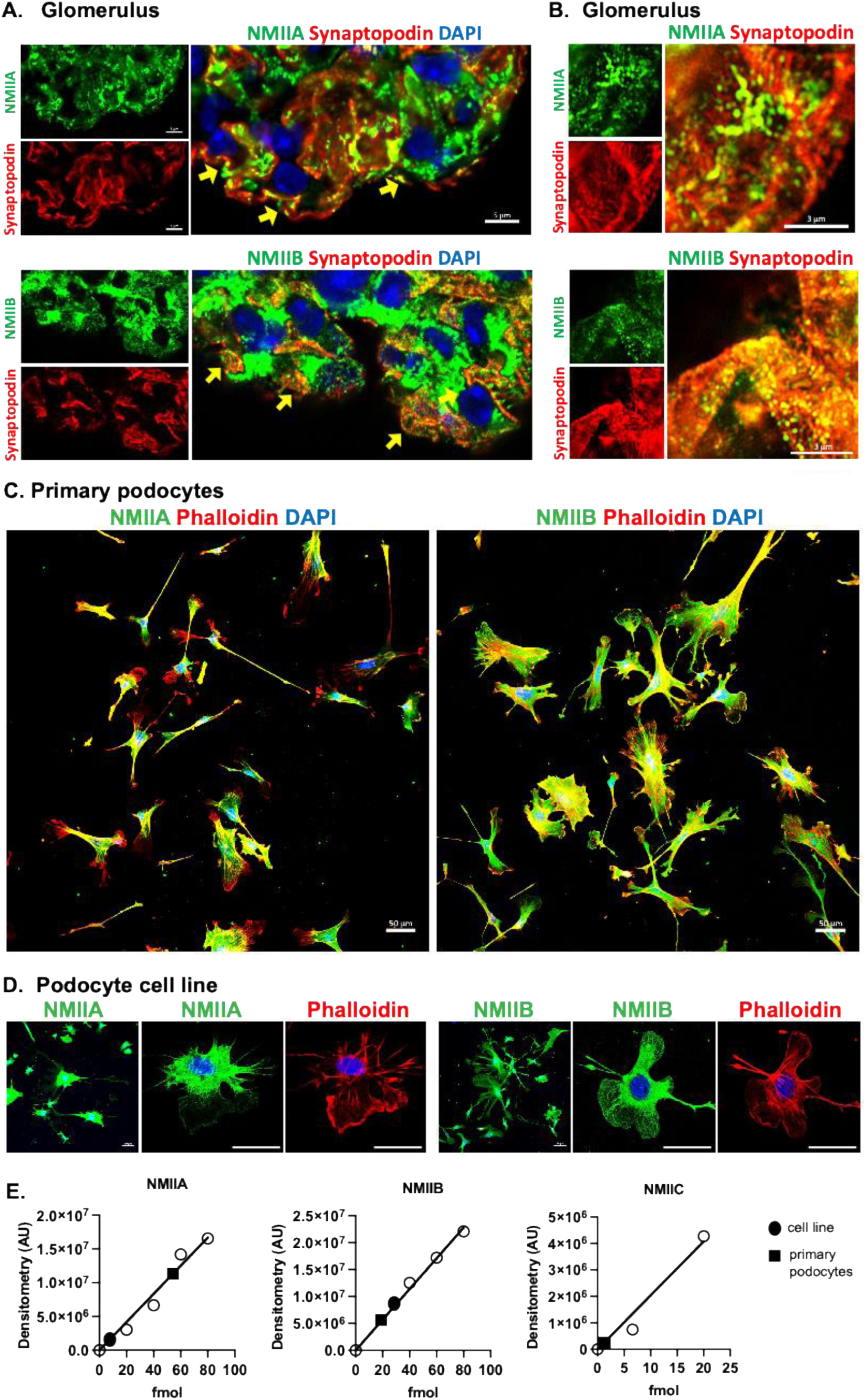
Distribution of myosin-IIA/IIB (NMIIA/NMIIB) in mouse glomeruli and isolated podocytes. A) Isolated mouse glomeruli were co-stained for the podocyte marker synaptopodin and NMIIA or NMIIB as indicated, to show co-localization in intact mouse glomerular podocytes *in situ.* The scale bar represents 5 μm. B) Super resolution microscopy of glomerular capillaries in cryosectioned kidney cortex show robust co-localization of NMIIB with synaptopodin positive podocytes. The scale bar represents 3 μm C) Isolated primary mouse glomerular podocytes show differential distribution of NMIIA and NMIIB, where NMIIA is localized centrally in majority of the cells and NMIIB is distributed broadly, including lamellipodial extensions. The scale bar represents 50 μm. D) Podocyte cell line developed from primary mouse glomerular podocyte outgrowths shows preservation of differential distribution of NMIIA and NMIIB. Scale bar represents 50 μm. E) Quantification of NMII paralog expression in equivalent amounts of lysates from a WT podocyte cell line (filled circle) and isolated primary podocytes (filled square) by comparison to standard curves generated using purified proteins (open circle). NMIIC was only detectable in primary podocytes.

Primary podocyte outgrowths stained for NMIIA and NMIIB showed distinct pattern of distribution, with NMIIB localized broadly to membrane edges and lamellipodial extensions while most of NMIIA was centrally located (Figure 1C). These images are consistent with a lesser degree of observed overlap of NMIIA signal with synaptopodin than NMIIB in glomeruli, suggesting that compared to NMIIA which is known to be more abundant than the other paralogs, a greater proportion of NMIIB is localized to dynamic membrane blebs and extensions. Preservation of differential distribution of NMIIA and NMIIB in a podocyte cell line developed from WT primary podocyte outgrowths support use of podocyte cell lines for studies related to WNK1 and actomyosin signaling (Figure 1D).

To determine whether the total amount of NMII paralogs expressed is a contributing factor for differential localization of NMIIA and NMIIB, we quantified the amount of the protein expressed in podocyte cell lines and primary podocytes by comparison to standard curves using purified proteins (Figure 1E). The calculated molar ratio of NMIIA/NMIIB was 0.17 in the WT podocyte cell line, and 3.0 in the primary podocyte. These values show that the primary podocytes have 17x more NMIIA than the developed WT cell line. NMIIC was undetectable in the podocyte cell line by immunoblotting even with 3x greater protein load. The amount of NMIIC/NMIIB in the primary podocytes was 0.03. Thus, the molar ratios of NMIIA:NMIIB:NMIIC is 3 : 1 : 0.03 in primary podocytes, and 0.17 : 1 : 0 in the WT podocyte cell line. The differences in amounts of NMIIA:NMIIB and the small amount of NMIIC in the primary podocytes may be from contaminating non-podocyte cells which account for <15% of the total cell number (Figure S5). In both cell preparations, NMIIB comprises a significant proportion of the total NMII paralogs in podocytes, and the expression of NMIIA and NMIIB are both comparable, not orders of magnitude different like it appears to be for NMIIC.

### WNK1 kinase activity affects localization and activation of NMIIs

To test the hypothesis that WNK1 activity affects localization of NMII in podocytes, we treated isolated glomeruli with the WNK1-specific inhibitor WNK-IN-11 (W11) [29], and compared the distribution of NMIIs. NMIIB co-localization with synaptopodin in the capillary loops was markedly reduced in many areas while NMIIA co-localization was not as affected (Figure 2A). These results suggest while NMIIA and NMIIB are both present in podocytes, NMIIB localization may be affected to a greater extent than NMIIA by WNK1 inhibition.

**Figure 2.**
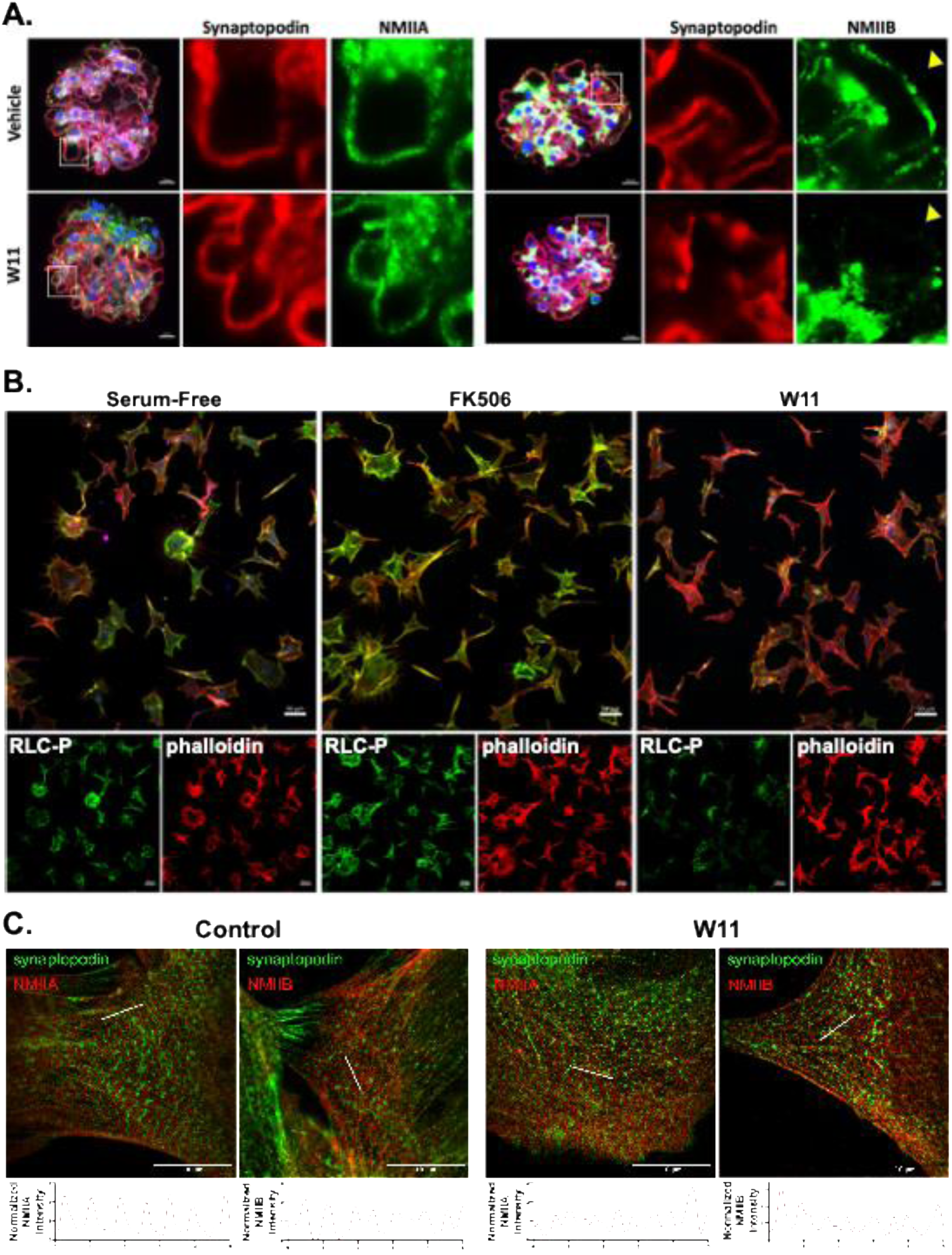
Acute effects of WNK1 inhibition on glomerular function and NMII paralog localization. A) Isolated mouse glomeruli after 2 hour treatment with control (vehicle) or 1 µM WNK1-specific inhibitor (W11), co-stained for NMIIA/IIB and synaptopodin. First column of images show merged images of a representative glomerulus, stained as indicated after treatments. Separated channels for the selected region (white box) is shown enlarged. With W11 treatment, NMIIB staining was markedly reduced at many of the glomerular capillary loops (yellow arrowhead), where synaptopodin and phalloidin staining were still present. The scale bar represents 10 μm. B) Podocytes in culture were stained for phosphorylated RLC (RLC-P) after 2 hour treatment with FK506 to activate WNK1 kinase or co-treatment with FK506 and W11 to inhibit WNK1 kinase. Enlarged merged images are shown with separate channels shown below. Scale bar represents 50 μm. C) Sarcomere-like structures were induced in WT podocytes with VRAD medium to induce pathologic remodeling processes. The effects of WNK1 inhibition with W11 treatment was assessed by comparison of NMII stain periodicity (lower graphs).

As NMIIs are ATPases that are activated by phosphorylation of RLC, we evaluated whether WNK1 inhibition not only affected the localization of NMIIs but also its activity. WT cultured podocytes were treated with the calcineurin inhibitor FK506, which we showed increased WNK1 kinase activity, or FK506 and WNK1-inhibitor W11 to test whether FK506-induced changes can be attributed to WNK1 activation. Staining of the cells for phosphorylated RLC after treatments showed FK506 modestly increased RLC-P over serum free conditions. Co-treatment with W11 markedly reduced RLC-P levels below that of serum free conditions (Figure 2B), indicating that in cultured podocytes, myosins have a high basal activity that is reduced by WNK1 inhibition. These results suggest WNK1 activity contributes to maintenance of baseline NMII activity.

NMIIA forms SLSs with alternating synaptopodin staining pattern in the podocytes of glomerular injury models and in cultured podocytes after VRAD medium-induced differentiation, which simulate injured podocytes[17,18]. We asked whether WNK1 contributed to the maintenance of sarcomere-like structures by comparing the distribution of NMIIA and NMIIB in response to W11 treatment. We found that podocytes appeared to have a more diffuse pattern of synaptopodin and NMIIA (Figure S2) and NMIIB (Figure S3) staining pattern in response to W11. NMII intensity pattern comparison shows disruption of sarcomere-like structure periodicity, indicating disruption of the NMII structures with WNK1 inhibition with W11 (Figure 2C).

### Podocyte injury phenotype in Alport Syndrome includes WNK1 and actomyosin protein expression differences

In the COL4α3 knockout (KO) Alport Syndrome model (C57BL6/J background), the disease progresses relatively slowly with mice succumbing from renal disease at around 8 months of age [30]. Thus, earlier time points, 2 – 4 months, represent early stages of renal disease, where injured podocytes are viable, and renal function is preserved. Comparison of glomerular cell outgrowths from 4 month old WT mice showed uniform podocyte cell shape and size (Figure S4A), but heterogeneity in size and cell number in the KO glomerular cell outgrowths (Figure S4B). KO glomeruli showed variable behavior with evidently fewer smaller podocytes migrating away from the explanted glomerulus earlier, and large hypertrophied cells migrating at a significantly slower rate than those from WT glomeruli (Figure S4B).

Cells that migrated from decapsulated glomeruli were positive for the podocyte-specific marker WT1 (Figure S5) confirming that they are podocytes (>85% of total). The observed differences in cell migration and heterogeneity in cell shape are consistent with known pathology of glomerular diseases, where injured podocytes are broadly distributed throughout the glomeruli in the renal cortex.

To extend our studies to biochemical characterization of WNK1-mediated signaling to NMIIs, we generated cell lines from WT and KO primary podocytes (Figure 4A), that retained differential morphological features (Figure 3). KO podocytes had fewer lamellipodial extensions and many appeared polygonal in shape with reduced structural complexity (Figure 3A).

**Figure 3.**
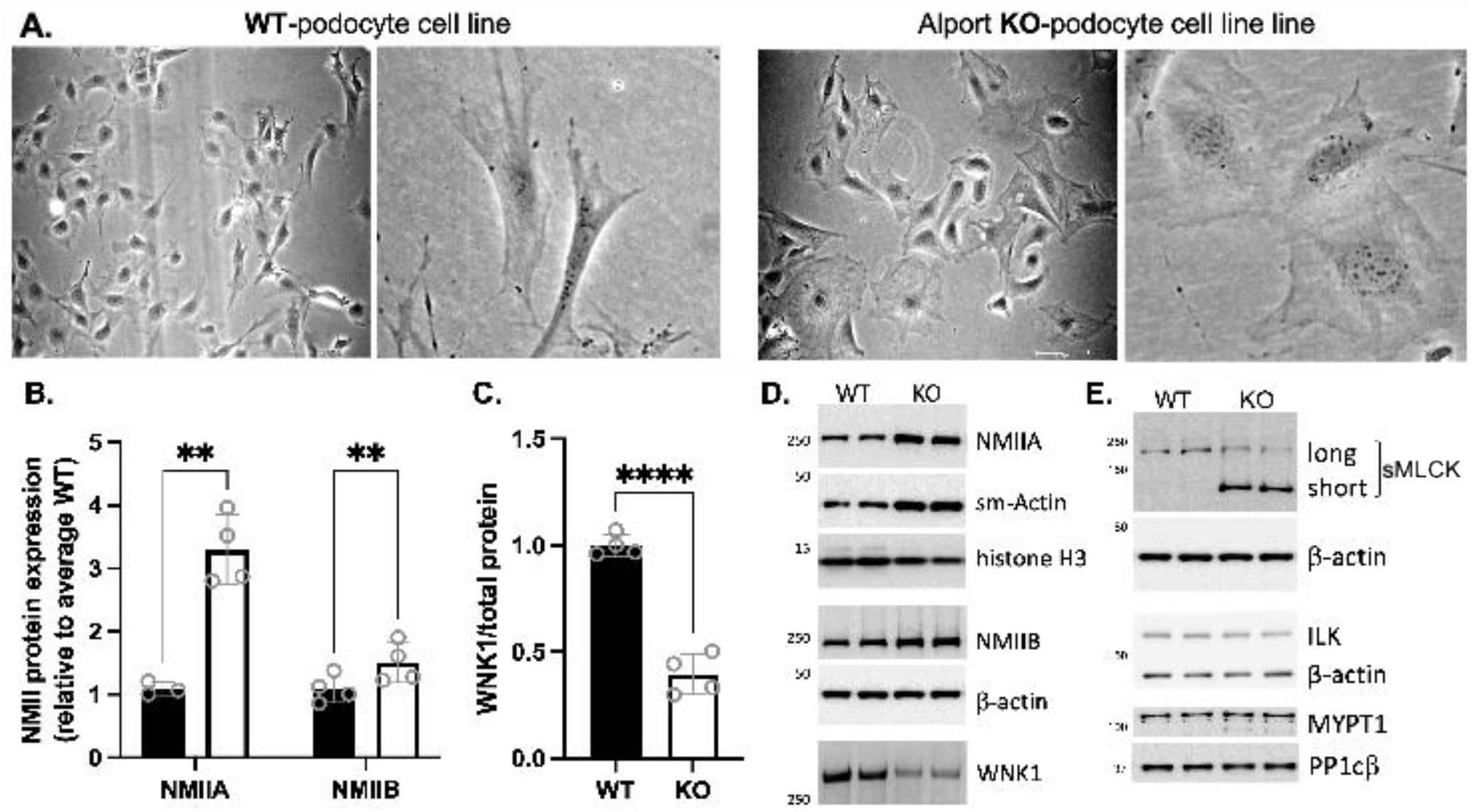
Characterization of podocyte cell lines developed from WT and *Col4a3^-/-^* Alport mouse glomeruli. A) Live cell microscopy of podocytes in culture. B) Comparison of NMIIA and NMIIB expression showed both were significantly increased in KO (white bar) over WT (black bar). C) WNK1 expression was significantly reduced in KO samples. Each dot represents average of triplicate experiments performed 3-4 distinct times and immunoblotted separately. Bars represent data normalized to mean of WT ± S.D. **P<0.01 by 2-way ANOVA, followed by Tukey’s multiple comparisons post-test, ****P<0.0001 by t-test, 2-tailed, using GraphPad Prism. D) Representative immunoblots of actomyosin proteins and WNK1. E) Representative immunoblots of selected regulators of NMII activity.

**Figure 4.**
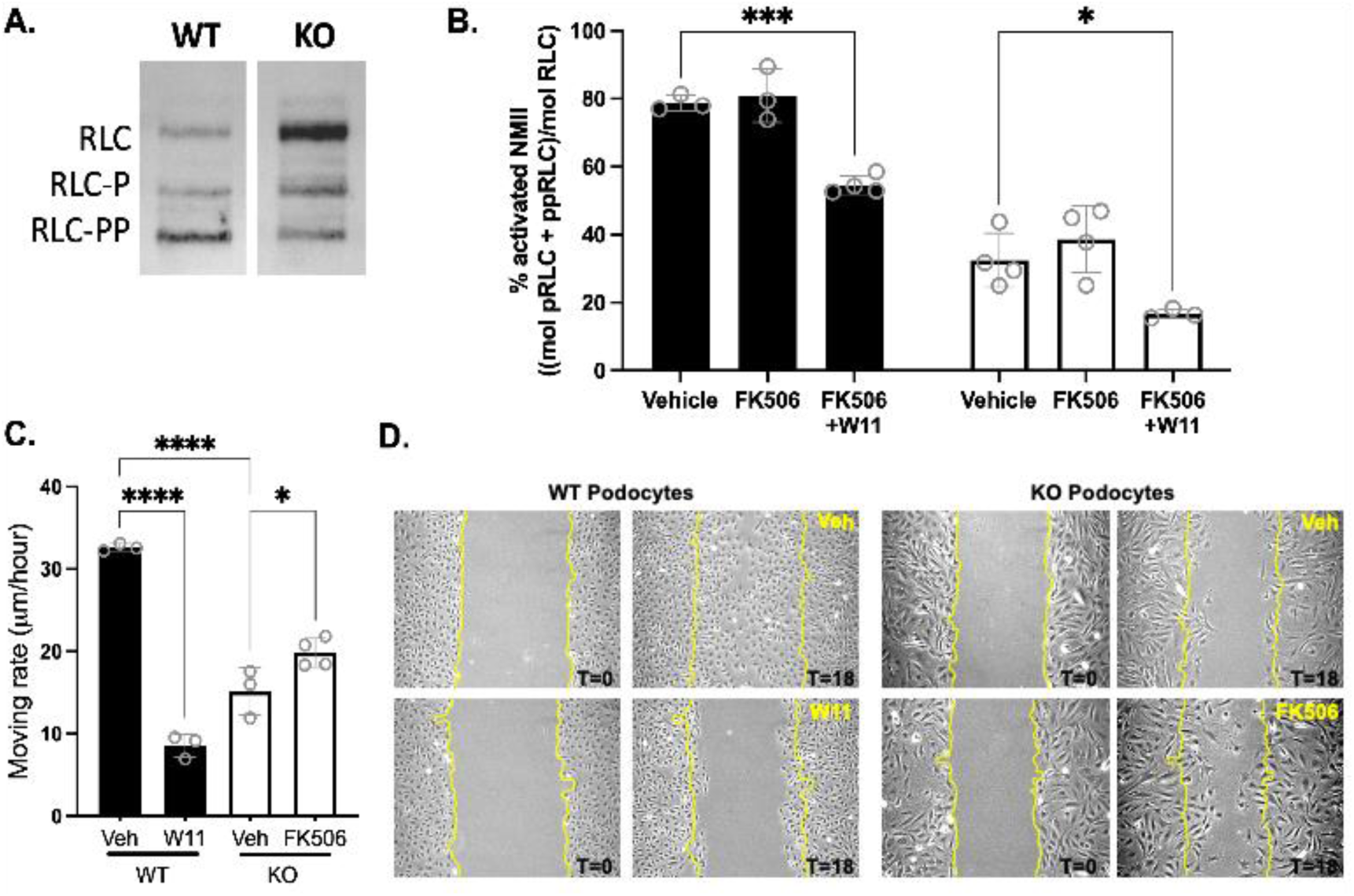
Effects of WNK1 activity on levels of activated NMII and cell migration. A) Representative immunoblot of WT and KO podocyte cell extract for myosin RLC after separation of mono-(RLC-P) and di-phosphorylated (RLC-PP) forms from non-phosphorylated RLC. Relative band intensities show that a greater amount of total RLC is di-phosphorylated (bottom band, RLC-PP) in WT podocytes, whereas a greater amount is unphosphorylated and inactive (top band, RLC) in KO. B) Graphed measurement of total activated NMII in WT and KO podocytes after indicated treatments, determined by calculated % of total RLC that is mono- or di-phosphorylated. Black bars are WT, white bars are KO. *P<0.05, ***P<0.001 by 2-way ANOVA, followed by Dunnett’s multiple comparisons post-test using GraphPad Prism. C) Effects of WNK1 activity on migration properties of WT and *Col4a3^-/-^* Alport KO podocytes. Comparison of quantified cell migration rates, as a measure of wound healing capability are shown. Dots represent average wound width measured in triplicate, from separate experiments; N=3 or 4. Significance was determined by ordinary one-way ANOVA, followed by Bonferroni’s multiple comparisons test using Graphpad Prizm; *P<0.05, ****P<0.0001. D) Representative images using the scratch wound-healing assay to induce podocyte migration. Cells were grown to confluency determined by dish surface area coverage. KO podocytes are larger and covered a greater surface area than WT podocytes (Figure 3A). Images were taken immediately after scratch wound (T0) and 18 hours after culture in normal FBS-supplemented media (T18), in the presence of WNK1 inhibitor or activator as indicated. WT podocytes subjected to WNK1-inhibition (W11), had a significantly reduced capacity to migrate into the wound area compared to vehicle control (Veh). Compared to WT at 18 hours, Alport KO-podocytes had a significantly reduced migration capacity under normal culture conditions (Veh). WNK1 activation with FK506 modestly increased the wound healing rate. The scale bar represents: 100 μm.

Based on the observed distinct morphology of WT and KO primary and cultured podocytes, we asked whether expression of WNK1, actomyosin proteins, and canonical regulators of NMII contractile activity were also different. Comparison of NMIIA and NMIIB expression in WT and KO podocytes showed both isoforms were significantly upregulated in KO cells, respectively by 3.3 ± 0.6 fold and 1.5 ± 0.3 fold (Figure 3B, D). Smooth muscle actin was also significantly increased in KO cells (2.4 ± 0.5 fold, N=4) compared to WT (Figure 3D), indicating activation of contractile protein expression. WNK1 protein was significantly reduced in the KO cells (0.4 ± 0.1 fold, N=4) compared to WT in both cell lines (Figure 3C, D) and primary cells from 2.5 month old mice (Figure S7).

Despite lack of significant lamellipodial features, more NMIIB along membrane edges was observed in KO podocytes (Figure S6B), where the specific substrate of WNK1 kinase, OSR1, was shown to be phosphorylated in response to FK506 [11]. These results support the hypothesis that WNK1 activity at podocyte membrane edges contribute to both NMIIA and NMIIB-mediated membrane dynamics.

The expression of selected regulators of NMII contractile activity, smMLCK and the subunits that form the MLCP, were compared between WT and KO podocytes (Figure 3E). WT podocytes expressed only the long form of smMLCK (Figure 3E), which is also known as non-muscle MLCK due to its expression in non-muscle cells. KO podocytes showed increased expression of the smooth muscle-specific short smMLCK isoform. The total smMLCK expressed (long + short form) was significantly higher in KO cells, which expressed a 3.7 ± 1.8 fold greater amount of total MLCK (Figure 3E). Interleukin-linked kinase (ILK), which is found in nascent and mature focal adhesions, can also phosphorylate NMII at its regulatory light chain (RLC) subunit, but the expression levels were comparable between WT and KO. Surprisingly, the protein levels of the ubiquitously expressed regulatory subunit of the canonical MLCP, MYPT1, and the catalytic subunit PP1cβ, were also comparable between WT and KO cells, despite increases in the substrate and kinase amounts. Thus, compared to WT, KO podocytes have increased smMLCK, smooth muscle actin, as well as total NMIIA and B. These measurements suggest changes in actomyosin protein expression occur very early in the disease process, and that greater substrate availability for smMLCK in the KO podocytes may be a compensatory mechanism to retain podocyte traction force.

### Activation of NMII is affected by WNK1 activity in WT and KO podocytes

Reduced colocalization of NMIIB and synaptopodin in podocytes on glomerular capillaries in response to WNK1 inhibition by W11 treatment (Figure 2A), suggests that NMIIB contractile activation is sensitive to WNK1-activity. NMII activation is dependent on the phosphorylation of its regulatory light chain (RLC) at Ser-19 (RLC-P) or Ser-19 and Thr-18 (RLC-PP) [31]. Based on increased NMIIs and smMLCK, we asked whether KO podocytes have more myosin activated by phosphorylation. To measure % of total activated NMII in podocyte cell extracts, we snap froze cells in 10% trichloroacetic acid by floating the dish in liquid nitrogen. Enzymatic reactions (phosphatases and proteases) are stopped in the conditions used, and thawed/precipitated cell proteins resolubilized/denatured in saturating urea conditions retain most phosphorylations. Phosphorylated RLCs were separated from non-phosphorylated RLC by glycerol gel electrophoresis and quantified to calculate the extent of phosphorylated RLC. Baseline comparison of activated NMII in WT and KO podocytes in culture showed that most of the NMII proteins in WT are activated by phosphorylation, while most of the NMII proteins in KO were inhibited (Figure 4A, B).

We measured the effect of activated WNK1 (FK506 treated), and inhibited WNK1 (W11 treated) on the extent of activated NMII (Figure 4B). Activation of WNK1 by FK506 did not induce a significant increase in total activated NMII as determined by the sum of total RLC mono- and di-phosphorylation, despite observable increases in Ser19 RLC-P by IF (Figure 2B). This discrepancy could be due to the high basal activation of myosins and the fact that NMIIs can be phosphorylated at its Thr18 or Ser19 residue. The glycerol gel separation does not distinguish between the two. Moreover, the total cell extract includes phosphorylated myosins that are localized to the membrane as well as all cytoplasmic myosins. Addition of W11 in addition to FK506 reduced the extent of activated NMII by about 20% in both WT and KO, where the total activated NMII was reduced from 81 ± 8% to 55 ± 3% in WT and from 39 ± 10 to 17 ± 1% in KO. These results suggest that WNK1 activity contributes to regulation of about 20% of total NMIIs in both WT and KO podocytes, presumably at the membrane edges of lamellipodial extensions where WNK1 appears to be most active[11].

Based on our immunoblot comparisons, KO podocytes have nearly 3-fold greater total NMII, and most of the myosins are in the inactive state, despite greater expression of total smMLCK (Figure 4A, B). We asked whether WNK1-activation by FK506 was sufficient to augment podocyte lamellipodia formation in injured podocytes despite a low level of activated NMII. Scratch wound healing assays using WT and KO podocytes showed that in the presence of FBS (Veh), WT podocytes could migrate to fully cover bare zones of the dish within 18 hours (Figure 4D). Inhibition of WNK1 with W11 over the same time frame, significantly reduced the migration rate of WT podocytes by over 3-fold from 33 ± 0.5 μm/hour to 8.5 ± 1.4 μm/hour, indicating that WNK1 activity is necessary for normal podocyte migration (Figure 4A, 4D), consistent with observations previously made using a pan-WNK inhibitor and WNK1 siRNA[11]. Compared to WT podocytes, KO podocytes had significantly reduced rate of migration under FBS supplemented conditions (Figure 4C, D), consistent with low extent of activated NMII (Figure 4A, B). Activation of WNK1 in KO podocytes using FK506, significantly augmented the migration rate from 15 ± 0.7 μm/hour to 20 ± 0.9 μm/hour (Figure 4C, D). These results collectively show that RLC-phosphorylation-dependent NMII activity is partly regulated by WNK1-activity. Thus, although the amount of NMIIA and NMIIB at the membrane cell extensions is a small proportion of the total, and the amount of WNK1 kinase was further reduced in the KO cells, increased WNK1 activity was sufficient to augment podocyte cell motility in injured podocytes.

### WNK1 kinase augments podocyte membrane extensions through activity-dependent signaling complexes

During glomerular disease in Alport Syndrome, podocyte foot processes efface[32], leading to detachment from the glomerular basement membrane, despite compensatory increases in sarcomere-like structures that contribute to adhesion to the basement membrane[17], and progression of renal disease. As shown above, although WNK1 protein is reduced in Alport KO podocytes (Figure 3C,D), cell migration is augmented by WNK1 activation that in turn increases NMII activity at the cell membrane (Figure 4C,D). Given the comparatively greater increase in NMIIA than NMIIB in the KO podocytes (Figure 3A), we asked whether the modest increase in NMIIB expression contributes to WNK1 kinase activity mediated effects on podocyte motility. Using immunofluorescence co-staining experiments, we found that WNK1, MYPT1, NMIIB and vinculin are enriched in and colocalize at undulating active membrane edges (Figure 5A). To test the hypothesis that these proteins physically interact, we performed co-IP experiments with anti-IgG control or anti-WNK1 antibodies and immunoblotted for the proteins indicated (Figure 5B). The results show that WNK1, NMIIB, MYPT1, and vinculin all selectively co-IP with WNK1 over control IgG.

**Figure 5.**
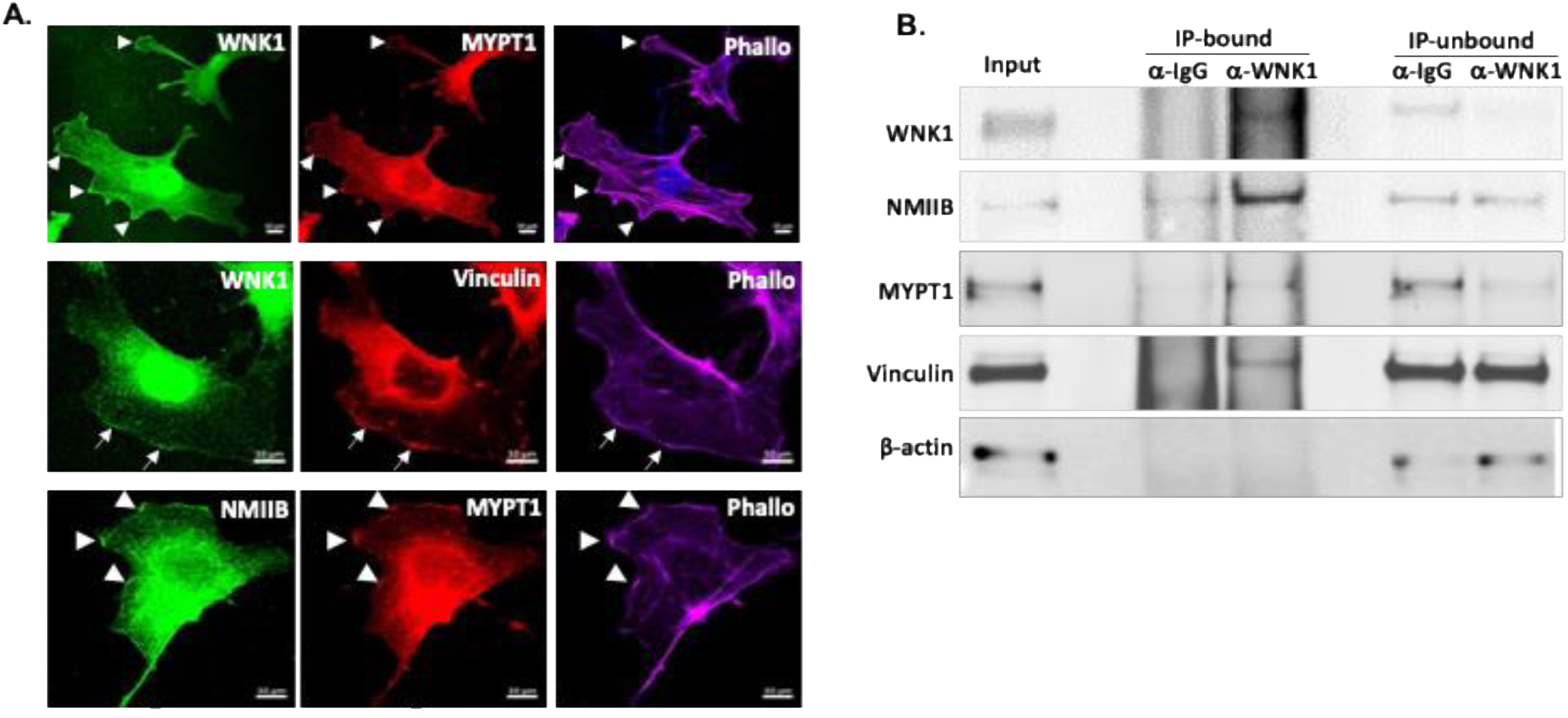
Evidence of WNK1 kinase interaction with actomyosin and focal adhesion proteins. A) Representative images of WT podocytes showing co-localization of WNK1 with MYPT1 (upper row), WNK1 with vinculin (middle row) and NMIIB with vinculin (bottom row). Arrowheads and arrows point to membrane undulations where WNK1 kinase appears to be enriched. The scale bar represents: 10µm. B) Immunoblot of selected proteins that co-immunoprecipitated with WNK1 antibody. Compared to IgG control, NMIIB, MYPT1, and Vinculin all co-IP selectively with WNK1 kinase. Reduction in the unbound fraction was seen for WNK1 and MYPT1, but not for NMIIB and Vinculin, consistent with co-localization studies in panel A that shows a small proportion of the total is found co-localized with WNK1 at the membrane edges. β-actin was used as a control for loading and indicator of IP-sample washing efficiency. Selected immunoblots from multiple co-IP experiments are shown.

Based on evidence of WNK1 kinase activity regulation by FK506 that increases cell motility (Figure 4C, D), and interaction with key focal adhesion proteins, NMIIB and vinculin (Figure 5B), we hypothesized that WNK1 activity is a determinant of podocyte membrane extension through signaling to NMIIB and vinculin at the cell membrane and by extension, foot processes.

Analysis of the podocyte cell membrane at leading lamellipodial extensions shows WNK1 kinase activity is required for the observed undulating pattern of WNK1 at the membrane edges (Figure 6A). Inhibition of WNK1 kinase with W11 caused a decrease in vinculin staining at lamellipodial edges where phosphorylated cortactin is found (Figure 6B). The WNK1 activity-dependent enrichment of vinculin at membrane edges was coincident with NMIIB localization (Figure 6C), which was also reduced with WNK1 kinase inhibition. These observations are consistent with co-localization and co-IP studies using podocytes in culture (Figure 5B), that showed evidence of WNK1 kinase interaction with NMIIB and vinculin.

**Figure 6.**
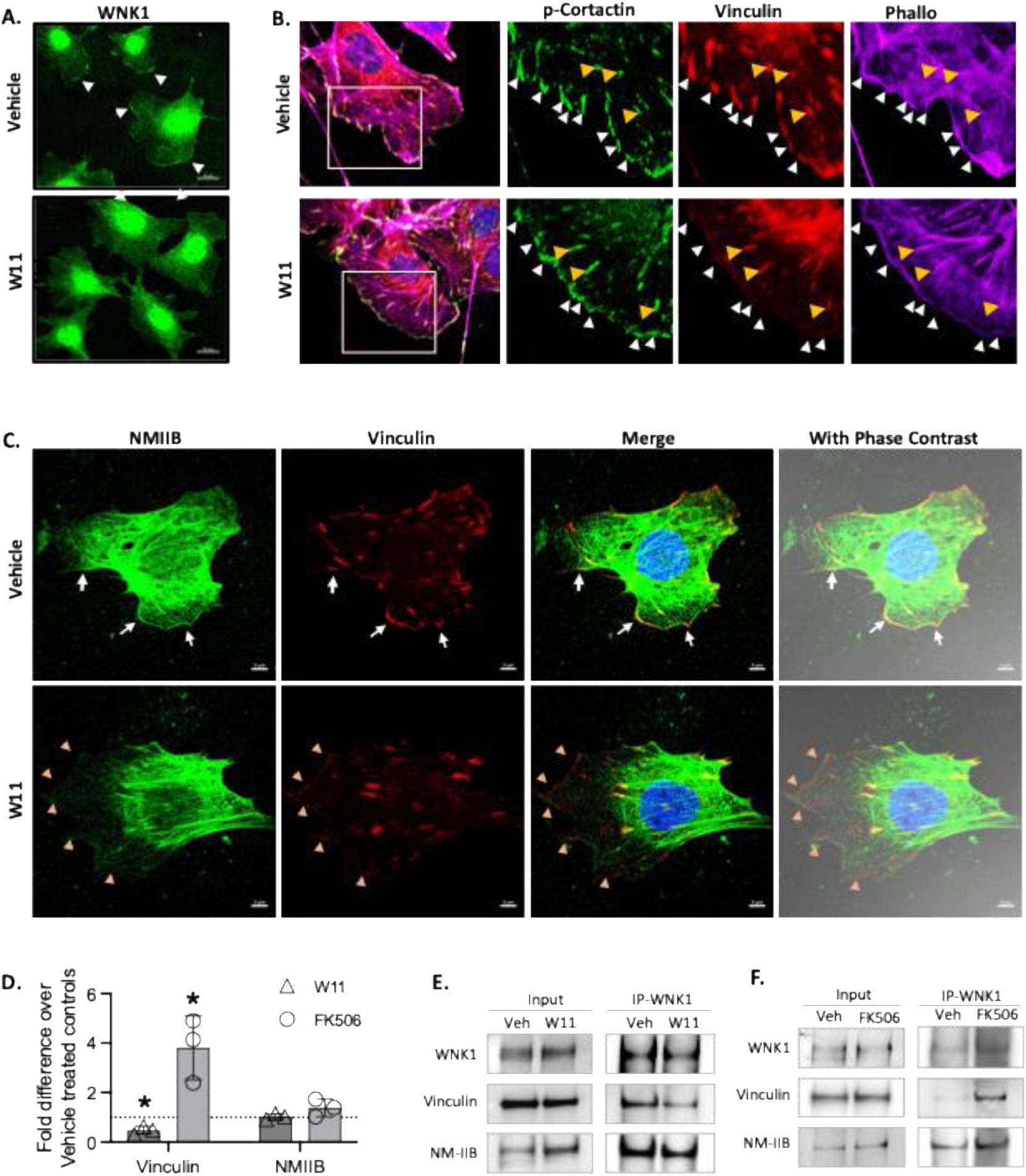
Membrane distribution of vinculin is decreased with WNK1 activity inhibition. A) WNK1 staining in podocytes shows WNK1 kinase localization to membrane edge undulations (arrowheads) was decreased from spread membrane with WNK1 kinase activity inhibition by W11 treatment. The scale bar represents: 20 µm. B) Effects of WNK1 kinase inhibition on co-localization of vinculin with p-cortactin (used as lamellipodia marker). Decreased vinculin from edge of lamellipodia (white arrows) but not cytoplasmic vinculin (yellow arrows) are indicated in enlarged images. C) Vinculin and NMIIB both decreased from leading edge of lamellipodia with WNK1 activity inhibition, but not in the cytoplasm where strong co-localization is evident in mature focal adhesions. White arrowheads in control points at leading edges of cells where undulating membrane edges show NMIIB colocalized with Vinculin. Yellow arrowheads in W11-treated cells point at the membrane edges with evident decreases in both NMIIB and Vinculin. The scale bar represents: 5µm. D) Comparison of vinculin and NMIIB protein co-immunoprecipitated with WNK1. *P<0.05 by T-test using Graphpad Prism. E) Representative immunoblots of NMIIB, Vinculin and WNK1 in WNK1-IP samples from podocytes treated with vehicle or WNK1-inhibitor (W11 1µM). F) Representative immunoblots of NMIIB, Vinculin and WNK1 in WNK1-IP samples from the podocytes treated with vehicle or WNK1-activitor (FK506 1µM).

To demonstrate that WNK1 kinase activity contributes to association of focal adhesion complex proteins, co-IP was performed using stimulated podocytes (10% FBS) under WNK1 inhibited (W11) or WNK1 activated conditions (FK506). Comparison of immunoprecipitated proteins showed that a greater proportion of vinculin co-IPed with activated WNK1 kinase than with inhibited WNK1 kinase (Figure 6D, E, F). More NMIIB co-IPed with activated WNK1 kinase, but the value was not statistically significant.

To extend the cultured cell findings to glomerular podocytes *in situ*, isolated glomeruli were treated with vehicle or a WNK1 kinase inhibitor and assessed for evidence of changes in localization of NMIIB and vinculin. Representative images of rendered Z-stack images in series (9 slices, 8um), show that NMIIB localization in podocytes of outer capillary loops in control glomeruli (arrows) is reduced after W11 treatment (Figure S8A). Glomeruli stained for vinculin show marked reduction in staining with WNK463 treatment (Figure S8B), consistent with results from podocytes in culture (Figure 6B, C). Collectively, these studies support the hypothesis that WNK1 kinase activity contributes to regulation of podocyte membrane dynamics through association with focal adhesion proteins, which include not only NMIIA but also activated NMIIB.

### WNK1 activity is required for maintenance of podocyte foot processes in glomeruli and *in vivo*

Based on work using podocyte cell lines that showed WNK1 kinase activity regulates membrane dynamics in lamellipodial extensions to affect actomyosin activity and localization of focal adhesion proteins like vinculin, we hypothesized that acute inhibition of WNK1 may affect the filtration function of glomeruli *in vivo*. As illustrated in Figure 7A, WT mice were treated with WNK463 by oral gavage (10 mg/kg), then injected with ½ N saline (i.p.) to increase urine flow. Urine was collected over 3 hours, after which kidneys were collected. WNK463 treatment significantly increased mouse urine Albumin/Creatinine ratio (ACR) from 0.1 to 0.6 mg/mg (Figure 7B,C) and the difference in urine albumin was visible by Coomassie staining of urinary proteins after SDS-PAGE. Cryosectioned kidney cortex stained for synaptopodin showed marked reduction in glomerular staining in mice that were fed WNK463, similar to isolated WNK463-treated glomeruli (*ex vivo*)[11]. These results show that diminished synaptopodin staining of the glomeruli corresponds to reduced permselecticity and may reflect podocyte foot process effacement.

**Figure 7.**
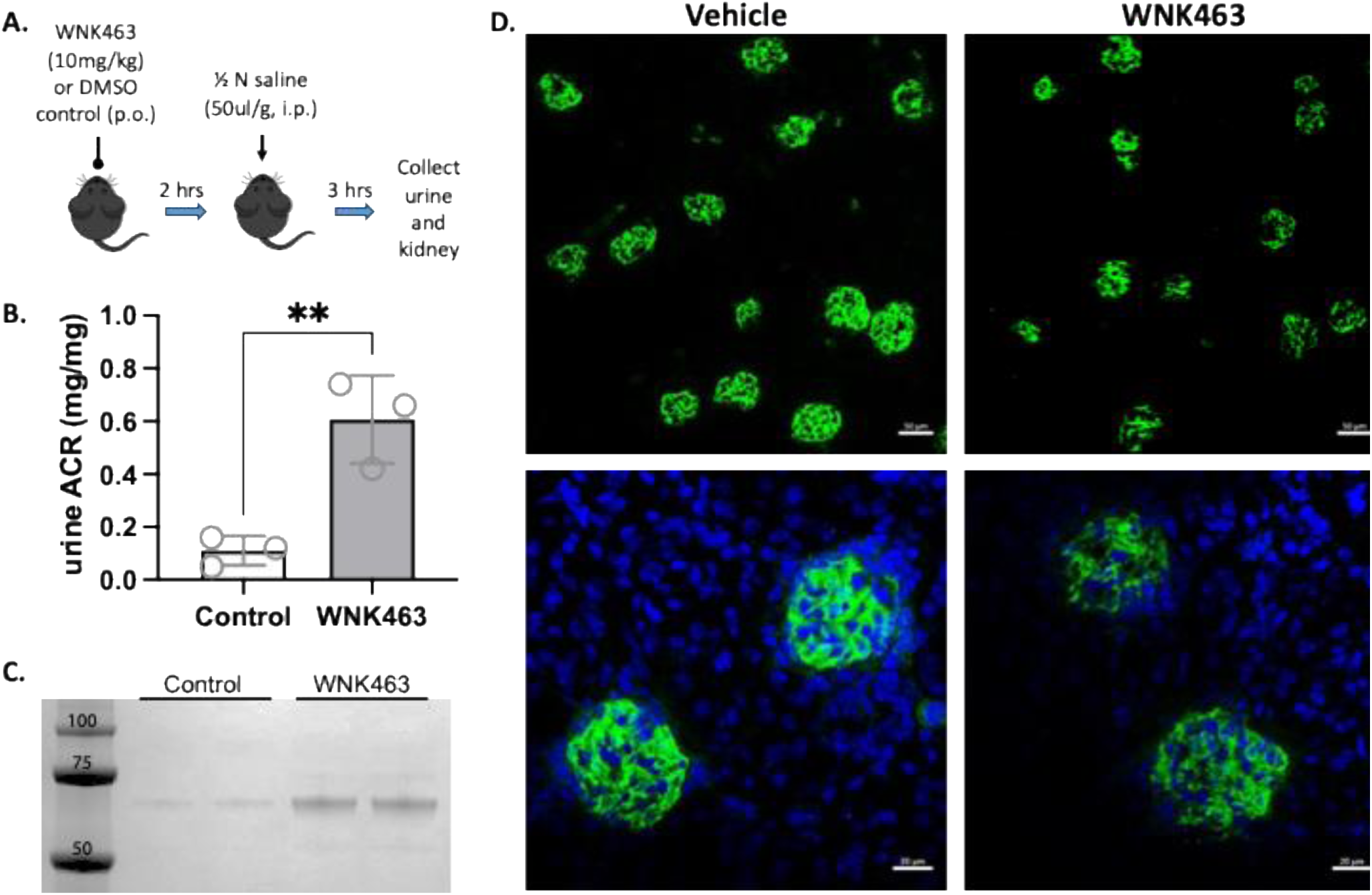
WNK1 kinase activity is required for maintenance of glomerular function. A) Schematic of *in vivo* acute inhibition of WNK1. WT mice were given WNK inhibitor or equivalent volume of DMSO vehicle by oral gavage (p.o.), 2 hours before i.p. injection of ½ N saline to increase urine output. Urine collected over a 3 hour period were pooled. Kidneys were collected at the end of 3 hours and embedded in OCT. B) Urine ACR, comparing mice treated with control or WNK463. C) Coomassie stained gel of urinary proteins separated by SDS-PAGE. Volume loaded were adjusted by creatinine values. D) Cryosectioned kidneys collected at end of study were stained for synaptopodin (green) and nuclei (DAPI, blue). Scale bar: 50 μm (upper row), 20 μm (lower row).

### WNK1 activation improves Alport KO mice glomerular structure

On the surface of capillaries, WT podocytes extend cell projections to form foot processes and slit diaphragms with complementary structures in neighboring podocytes. Alport KO podocytes have reduced cell projections, resulting in foot process effacement. Based on evidence of WNK1 activity-mediated effects on synaptopodin localization, which coincide with WNK1, NMIIB, and vinculin presence at dynamic membrane edges in cultured podocytes, we asked whether podocyte structure in the KO glomeruli could be improved with WNK1 activation. We hypothesized that FK506 could contribute to retention of podocytes during Alport disease progression by restoring podocyte structure including foot processes, thereby enhancing the attachment of podocytes to the glomerular basement membrane. Staining of podocyte-specific synaptopodin after treatment with vehicle or FK506 in combination with WNK463, showed that as previously published, WNK1 activity was required for continuity of glomerular capillary synaptopodin staining (Figure 8A, upper row). Glomeruli from Alport KO animals with decreased podocyte number have discontinuous and reduced synaptopodin staining that could be improved with acute FK506 treatment. WNK463 blocked the improvement associated with FK506 treatment and further diminished the intensity of synaptopodin staining, suggesting it further disrupted the glomerular structure (Figure 8A, lower row).

**Figure 8.**
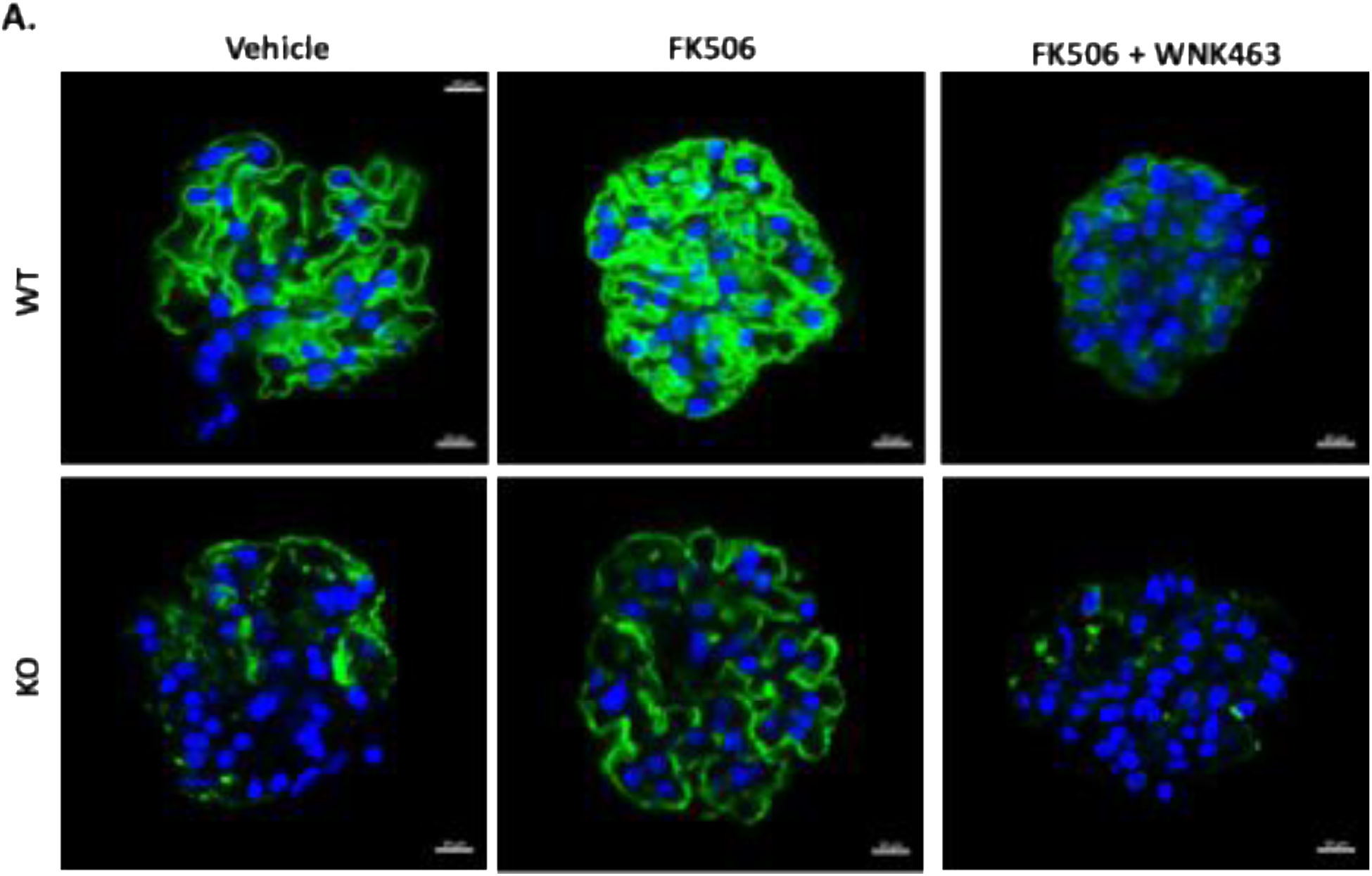
WNK1 kinase signaling directly regulates podocyte foot process structure in situ. A) Representative synaptopodin (green) and DAPI (blue) stained glomeruli from WT (upper row) and Alport KO (lower row) mice after *ex vivo* treatments as indicated.

These results collectively shows that WNK1 kinase activity is necessary for normal podocyte cytoskeletal and membrane dynamics and that disruption of WNK1 kinase activity leads to loss of activated actomyosin and nascent focal adhesions at the leading lamellipodial edges of podocytes. Perturbation of WNK1 kinase activity causes structural changes in glomerular podocytes *in situ*, resulting in reduced synaptopodin staining in isolated glomeruli (Figure 8, upper row), and reduced myosin activation in podocytes (Figure 2B). Acute WNK1 kinase inhibition *in vivo* recapitulates the podocyte structural and physiological phenotype of Alport KO mice with proteinuria (Figure 7). Activation of WNK1 in glomeruli from Alport KO mice showed partially restored synaptopodin staining, suggesting: 1) decreased synaptopodin staining in KO glomeruli may reflect structural perturbations including foot process effacement, and 2) the structure of injured podocytes from Alport KO animals may be improved by WNK1 activation by FK506. These observations are consistent with studies where FK506 treated patients with glomerular diseases had improved disease course and proposes a mechanism of action that includes WNK1 activation.

## Discussion

In several studies, CNIs, cyclosporin A and tacrolimus (FK506), reduced proteinuria in patients with nephrotic syndrome [33–39]. Mouse models of nephrotic syndrome suggest that CNIs exert their podocyte protective effects through suppression of inflammatory responses as well as non-immune mediated mechanisms of actin including synaptopodin stabilization[40,41]. However, the reported effects of CNIs on podocytes were empirical in nature without definition of causal mechanisms. Our studies present WNK1 kinase activity-dependent maintenance of podocyte foot processes as a potential mechanism involving vinculin and actomyosin activity that can be acutely and directly manipulated by CNIs.

Using podocyte cell lines, primary podocytes, as well as isolated glomeruli, we found differential distribution of NMIIA and NMIIB in healthy and diseased podocytes from an Alport model. Our novel discovery of a distinct pool of NMIIB present at the edge of lamellipodial extensions, and evidence of a WNK1-NMIIB signaling axis in glomerular podocytes supports the hypothesis that WNK1 activity contributes to the regulation of cell motility and membrane extensions, in part, by signaling not only to NMIIA but also NMIIB.

Early manuscripts describing podocytes in culture showed different podocyte cell shapes that were attributed to culture conditions[42,43]. We found that WT and Alport KO primary podocytes under matched conditions have distinct cell shapes that are similar to those found in our conditionally immortalized WT and KO cell lines that were expanded in the presence of interferon at 33°C, and differentiated in 37°C. These differences in podocyte structure and behavior represent the effects of the different biologic processes associated with normal versus Alport syndrome environments. Persistence of differences in culture after the cells leave the glomerular capillaries indicates that the KO podocytes have intrinsic differences from WT cells that are characteristic of their disease state and independent of external factors such as interactions with other cells, the glomerular basement membrane, or culture conditions.

In non-muscle cells, NMII activity is regulated by phosphorylation of its RLC subunit. Re-distribution of the NMII fluorescence signal in response to WNK1 inhibition in intact glomeruli suggests that the myosin network is in part regulated by WNK1 kinase mediated activation of MLCK or inhibition of MLCP. MLCP activity inferred from measurements of phosphorylated MYPT1/total MYPT1 in podocytes treated with W11, were not significantly different, suggesting reduction in activated myosin in response to W11 is attributed to reduced phosphorylation of RLC. Several kinases have been shown to phosphorylate RLC in non-muscle cells, including ILK, DAPKs, ROCK, as well as the canonical MLCK. Notably, two splice-variants of MLCK are expressed in primary podocytes and podocyte cell lines, the long non-muscle MLCK and the shorter smooth-muscle MLCK[21]. While in cultured cell lines, long MLCK is often exclusively expressed[22], we discovered that in KO podocytes, the short smooth muscle MLCK is the dominant variant. This was an unexpected finding, given that most contractile smooth muscle cells rapidly de-differentiate in culture, cease to express the shorter smooth muscle MLCK, and re-express the long variant. Long and short MLCK have comparable catalytic rates[22], and the region of the kinase that is expressed in the long form partly contributes to localization of the kinase during mitosis [44]. RNAseq analysis of differentially expressed genes in primary podocytes from WT and Alport KO mice showed over-representation of genes associated with the Lysosome, Focal Adhesion, and Cell-Substrate Junction (Figure S9) and increased expression of MRTFa/b regulated genes that are associated with myofibroblast differentiation[45]: TAGLN (2.5-fold increase in KO), ACTA2 (9.5-fold increase in KO), consistent with the KO podocyte cell line that showed increased smooth muscle actin expression (Figure 3D). Whether re-expression of the short smooth muscle MLCK in disease-model podocytes reflects compensatory responses to alter shape or to retain strong cell-cell or matrix attachments, and whether this change represents a broader change in gene expression profile remains to be investigated.

NMIIs have two phosphorylation sites on the RLC that regulate myosin ATPase activity. The physiological significance of di-phosphorylated RLC is unclear, given that it does not augment mono-phosphorylated myosin ATPase activity and is not found at significant levels in activated smooth muscle tissue. Several mechanisms of NMII regulation in podocytes have been described by others [46–48] that act through known regulators of NMIIs, MLCK and ROCK. MLCK and ROCK differentially activate peripheral and central pools of NMIIs in cultured cells to respectively promote mono- and di-phosphorylation of RLC[49]. This pattern is reminiscent of the central distribution of NMIIA and peripheral or extension-associated distribution of NMIIB in primary and cultured WT and KO podocytes (Figure 1, S6).

Pathologic remodeling of podocytes include increased expression of NMIIA (Figure 3A), and the appearance of SLSs patterning with synaptopodin *in situ*[17], which can be replicated in both primary podocytes and in podocyte cell line after differentiation in VRAD-media) [18]. Interestingly, we found that NMIIB also form SLSs like NMIIA. A modest increase in NMIIB that we observed (Figure 3A), may have been undetected in prior immunofluorescence imaging by others due to differences in fixation methods[16], and obscuring of low intensity signal in the podocytes, by higher intensity signals from endothelial cells that express greater amounts of NMIIB, necessitating focused imaging of capillaries (Figure 2A, S8). The collective evidence in the current work augments the importance of both NMIIA and NMIIB, which has received relatively little attention in studies of podocyte cell biology.

NMIIA and NMIIB are both expressed in glomerular podocytes and based on comparative studies of podocyte cell morphology and function, they are important determinants glomerular stiffness, a characteristic that reflects glomerular capillary and podocyte integrity[1,11]. Moreover, WNK1 contributes to regulation of vinculin and NMIIs to affect podocyte cell migration and contractility, reflective of podocyte membrane and actomyosin dynamics *in situ*, on glomerular capillaries. The disruption of SLS patterning after inhibiting WNK1, further supports a link between WNK1 pathway and NMIIs in pathological remodeling of injured podocytes. The cytoskeletal proteins, myosins and the regulated signaling by WNK1, MLCK and MLCP involved in membrane dynamics *in vitro* may reflect the NMIIs, actin and foot process structures *in vivo*. Thus, our findings suggest that reported improved GFR in response to CNIs may be partly attributed to NMII activity and membrane WNK1-NMIIB signaling axis in podocytes[32], and NMIIB activation may be targetable in early stages of podocyte injury to delay or reverse foot process effacement and podocyte loss in glomerular diseases.

## Materials and Methods

### Isolation of Mouse Glomeruli

To isolate glomeruli, kidneys were removed from mice, decapsulated, and dissected on ice to separate the cortices from the medullae. The cortices were minced with a razor blade and pushed through a screen (180 μm, W.S. Tyler Co, Cleveland, OH, United States) with a flexible metal spatula. The minced tissue was suspended in DPBS with 5.5 mM glucose and 0.3 mM pyruvate (GIBCO), filtered through a 90 μm nylon mesh (Falcon) to remove vessels and large pieces of tissue. The filtrate was collected in a 45 μm mesh nylon filter (Falcon) and contained intact glomeruli with minimal contamination. Glomeruli were maintained in DMEM with 0.1% FBS at room temperature and treated with drugs for immunostaining or seeded onto collagen I-coated dishes for 4-6 days to allow primary glomerular cell outgrowth. Primary podocytes were then trypsinized and filtered with 20 μm cell strainer, and re-plated for experiments.

### Ethical Approval

Animal research was performed in accordance with the UT Southwestern Medical Center Animal IACUC guidelines. The research study protocol (number 2014–0078) was approved by the UT Southwestern Medical Center Animal IACUC (NIH OLAW Assurance Number A3472-01). UT Southwestern Medical Center is fully accredited by the American Association for the Assessment and Accreditation of Laboratory Care, International (AAALAC). Animals are housed and maintained in accordance with the applicable portions of the Animal Welfare Act and the Guide for the Care and Use of Laboratory Animals. Veterinary care is under the direction of a full-time veterinarian boarded by the American College of Laboratory Animal Medicine. Mice were sacrificed for experiments by first anesthetizing with Avertin and then euthanizing by cervical dislocation.

### Immunofluorescence staining of cells and glomeruli

Cultured podocyte cell line or primary podocytes were reseeded onto collagen I-coated glass coverslips and allowed to attach for 24 hours. Cells on glass coverslips were then fixed for 10 minutes at room temperature with 4% paraformaldehyde in PBS, blocked with 3% BSA in PBS for 30 minutes, then permeabilized with 0.2% Triton-100 for 5 min. Staining was accomplished by incubating with primary antibodies overnight at 4°C, and then washing three times in PBS before incubation with secondary antibodies for 1 hour at room temperature. At the end of the incubations, samples were mounted onto glass slides using mounting medium containing 4′,6-diamidino-2-phenylindole (DAPI). For glomeruli staining, isolated floating glomeruli were maintained in DMEM with 0.1% FBS at room temperature and treated with drugs for 2 hours (W11 (1 μM), spun down at 500 g for 5 min, and then washed 3x in PBS. Washed glomeruli were resuspended in PBS, and pipetted onto a poly-prep slide. After glomeruli attached, the slide was rinsed in PBS to remove unattached glomeruli and further fixed with 4% paraformaldehyde for 20 min at room temperature. Fixed glomeruli on slide were then washed 3x in PBS, blocked with SEA BLOCK Blocking Buffer (PIERCE) for 1h, then permeabilized with 0.5% Triton X-100 in PBS for 5 min. Glomeruli on slide were then stained using standard procedures.

### Microscopy and Image Analysis

Confocal imaging was performed in the Cell Biology and Imaging Core in the O’Brien Kidney Research Core, on a Zeiss LSM880 with Airyscan laser scanning microscope equipped with Plan-Apochromat 10x/0.3 NA, 20x/0.8 NA, 25x/0.8 NA, and 63x/1.40 NA oil-immersion objective (ZEISS, Oberkochen, Germany). Fluorescence images were acquired using ZEN black 2.3 software with a 20x/0.8 NA or 63x/1.40 NA objective and Zeiss Immersion Oil 518F was used for the 63x/1.40 NA objective. Experiments were performed at constant room temperature. Images or regions of interest (ROIs) were further processed with ZEN 2.6 (blue edition) software.

### Scratch-Induced Cell Migration Assay

Equal numbers of podocytes were seeded on collagen-coated six well plates. Confluent cell monolayers were washed, and scratch wounds were created with a 1 ml pipette tip. Culture medium was replaced with culture medium containing DMSO, W11 (1μM), or FK506 (1μM). Images were obtained immediately after wound creation and before cell migration. Plates were returned to the incubator and cells were allowed to migrate for 18 h, at which time repeat images were obtained. Cell migration rates were calculated from three different regions of the wound width. Average of 10 independent measurements were tracked after 18 hours, and decreased wound width/hour was used as an estimate of cell migration rate.

### Immunoblotting

Cultured podocytes after indicated treatments, were gently rinsed in PBS, then overlayed with 1 mL 10% TCA/10mM DTT and snap-frozen by floating the culture dish on liquid nitrogen. After thawing on bench, cell precipitates were scraped off and collected in a microcentrifuge tube, collected by centrifuging for 1 minute at 1000 x g. Collected protein precipitates were aspirated free of TCA solution, washed in ethyl ether 3 x 15 minutes each, then air-dried in the chemical fume hood. Dried protein precipitates were completely solubilized in equivalent volume of urea sample buffer containing 8 M urea, 20 mM Tris (pH 8.6), 23 mM glycine, 10 mM DTT, 4 mM EDTA, and 5% sucrose. Protein concentrations were quantified by Bradford assay (Bio-Rad). Calculated amounts of proteins were then added to 0.25 volumes of 4xLDS sample buffer and reducing agent for SDS-PAGE per reagent instructions (Thermo Fisher). Protein solubilization quality and gel loading for immunoblots were predetermined by Coomassie-stained (Imperial Protein Stain, ThermoFisher) gel image analysis after separation by 4-12% SDS-PAGE (Bolt, MOPS buffer system, ThermoFisher). Separated proteins (2 to 8 μg per lane, depending on protein of interest) were transferred onto a nitrocellulose membrane (Biorad), then processed for immunoblotting using standard procedures. Briefly, membranes were incubated with specific antibodies diluted in 5% BSA/TBST overnight at 4°C, washed in TBST 3 × 10 minutes, probed with horse-radish peroxidase-conjugated secondary antibodies at room temperature for 1 hour, washed in TBST 3 × 10 minutes, then developed using ECL Plus (Thermo Fisher), and imaged using Chemidoc MP (Bio-Rad). The amount of protein to load per lane of gel was empirically pre-determined by a loading curve comparison for each antibody used.

### Co-immunoprecipitation studies for protein interaction

Podocyte were cultured on 10 cm collagen coated dishes to ∼80% confluency, then lysed in RIPA buffer with protease inhibitor (Halt protease inhibitor cocktail, Thermo). Lysates collected in microcentrifuge tubes were pelleted at 16,000 rpm for 1 minute, and collected supernatant fractions were equally divided into two for IP with equal amount of IgG and WNK1 antibodies (control-IP and WNK1-IP). To test the effects of WNK1 kinase activity inhibition or activation on protein interactions, cells were pre-treated with WNK1-inhibitor (W11, 1µM) in culture medium for 3 hours, or treated with WNK1-activitor (FK506, 1µM) in serum-free medium for 3hrs. Protein G beads (ThermoFisher) were used for immunoprecipitation following manufacturer’s instructions. Immunoprecipitated proteins were denatured in 1xLDS sample buffer with Reducing Solution (ThermoFisher), heated for 10 minutes, then separated in 3-8% Bolt mini-gels in a Tris-Acetate SDS Running Buffer system (ThermoFisher). Separated proteins were wet-transferred to 0.2 μm pore size nitrocellulose membranes (Whatman PLC, Buckinghamshire, UK) using a plate electrode Bolt mini-transfer system (ThermoFisher), then immunoblotted with specific antibodies using standard WB procedures.

### Measurement of RLC phosphorylation

RLC phosphorylation was measured by urea/glycerol-PAGE as previously described [50]. Podocytes snap-frozen in TCA and proteins processed in urea sample buffer as detailed above, were subjected to urea/glycerol-PAGE to separate mono- and di-phosphorylated RLC from non-phosphorylated RLC. Following electrophoresis, proteins were transferred to nitrocellulose membranes, and immunoblotted for pan smooth/non-muscle RLC. The molar ratio of mono- or di-phosphorylated RLC to total RLC was determined by quantitative densitometry of developed immunoblots and expressed as mol phosphate per mol protein.

### Podocyte cell culture

Podocyte cell lines were derived from WT and KO primary podocyte outgrowths from glomeruli isolated from adult mice (12-16 weeks of age). Primary podocytes were conditionally immortalized by infection with a lentivirus expressing the temperature sensitive SV40 T antigen tsA58. Lentivirus was produced from pLenti-SV40-T-tsA58 (abm cat# LV629) as previously described [51] using the VSV envelope from pMD2.G (Addgene cat#12259, gift from Didier Trono) and the packaging plasmid pCMV delta R8.2 (Addgene cat # 12263, gift from Didier Trono). Primary glomerular outgrowths were infected for 48 hours followed by selection with puromycin (2ug/ml). Conditionally-immortalized mouse podocytes were cultured in RPMI 1640 with 10% FBS. Cells cultures were expanded at 33°C in the presence of interferon, and for experiments, differentiated at 37°C for 7-10 days as previously described [52]. For imaging SLSs, WT podocytes were differentiated in VRAD medium in 37°C for 7 days, following a previously described method[53]. Briefly, the differentiation medium was supplemented with Vitamin D3 and retinoic acid to enhance podocyte-specific gene expression and promote the organization of sarcomere-like structures, thereby mimicking the phenotype of primary podocytes.

## Data availability statement

The data that support the findings of this study are available from the corresponding author upon reasonable request.

## Author contributions

ZL and ANC conceived and designed the research; ZL, EL, SJ, FA, JY, and ANC performed the research and acquired the data; ZL, SJ and ANC analyzed and interpreted the data; MAR, WPR, LAB and RTM provided critical reagents; ZL and ANC drafted the manuscript; ZL, ANC, LAB, HS and RTM contributed to revising the manuscript.

