## Supplemental Material for "WNK1 kinase activity is required for maintenance of podocyte foot process structure"

**Supplementary Material**

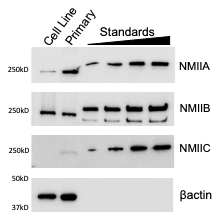

**Figure S1. Comparison of NMII paralog expression in WT podocyte cell line and primary podocytes isolated from WT mice.**  Podocyte cell lysates were quantified by Bradford assay and equivalent amounts loaded next to protein standards in the same gel to quantify molar ratios of specific paralog expression. For NMIIA and NMIIB, 1.5 ug or total protein were loaded. For NMIIC, 4.5 ug of total protein was loaded. Purified NMIIs were loaded from 20 to 80 fmol for NMIIA, NMIIB, and 7 to 42 fmol for NMIIC. Protein standards vary in size due to presence or absence of Halo tag. βactin was used as loading control for the podocyte cell lysates.

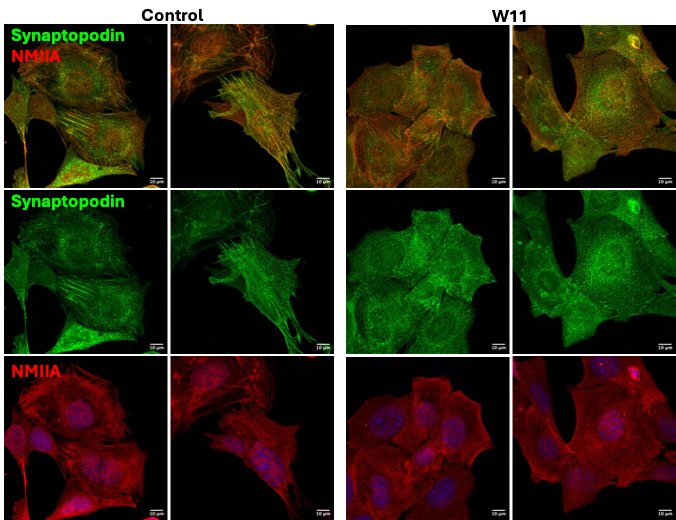

**Figure S2. Effect of W11 on the distribution of synaptopodin and NMIIA in VRAD-differentiated podocytes with sarcomere-like structures.**

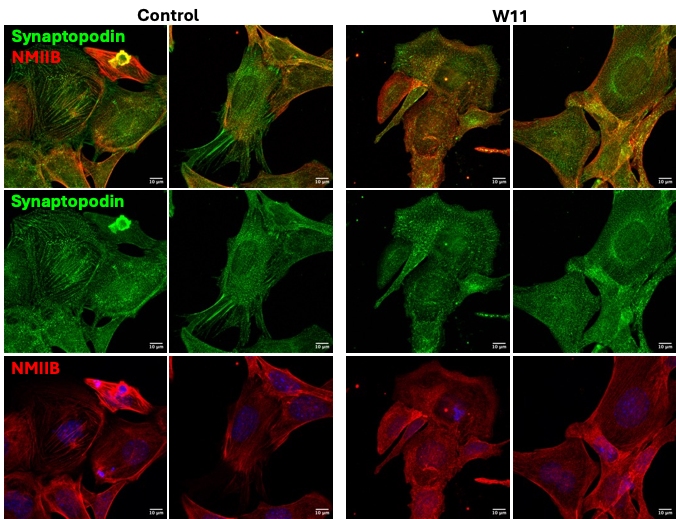

**Figure S3. Effect of W11 on the distribution of synaptopodin and NMIIA in VRAD-differentiated podocytes with sarcomere-like structures.**

**A.**

| 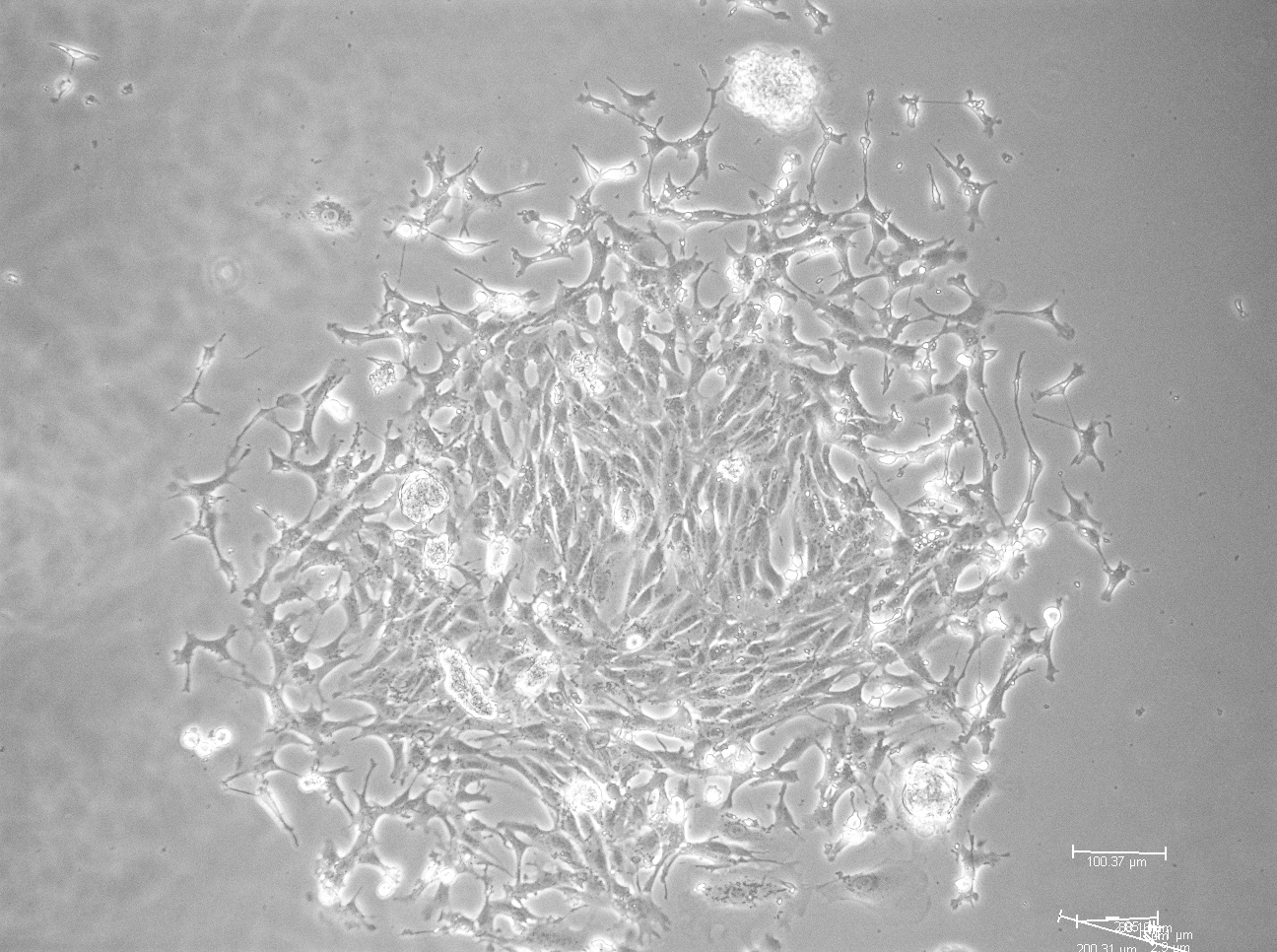 | 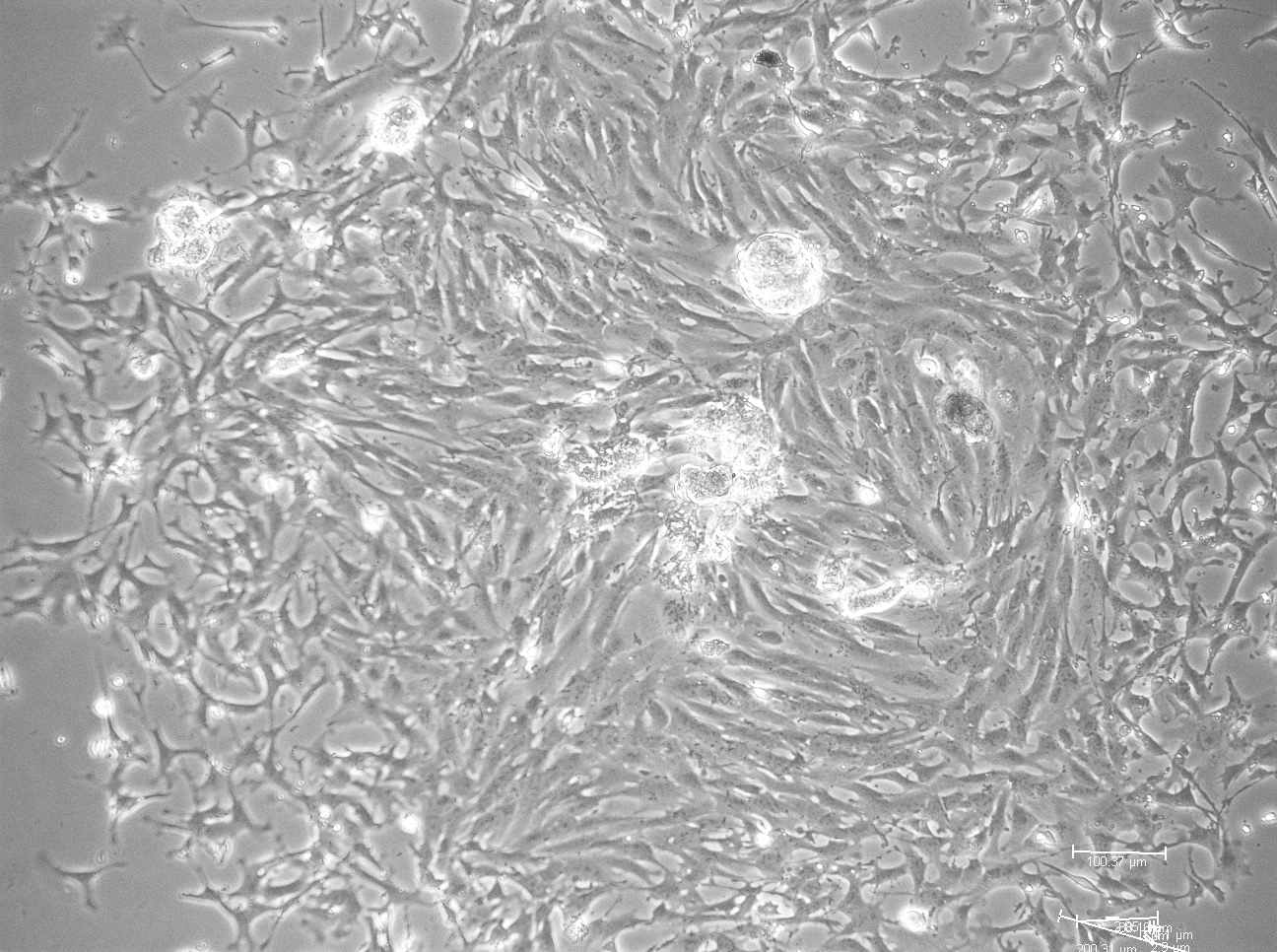 | 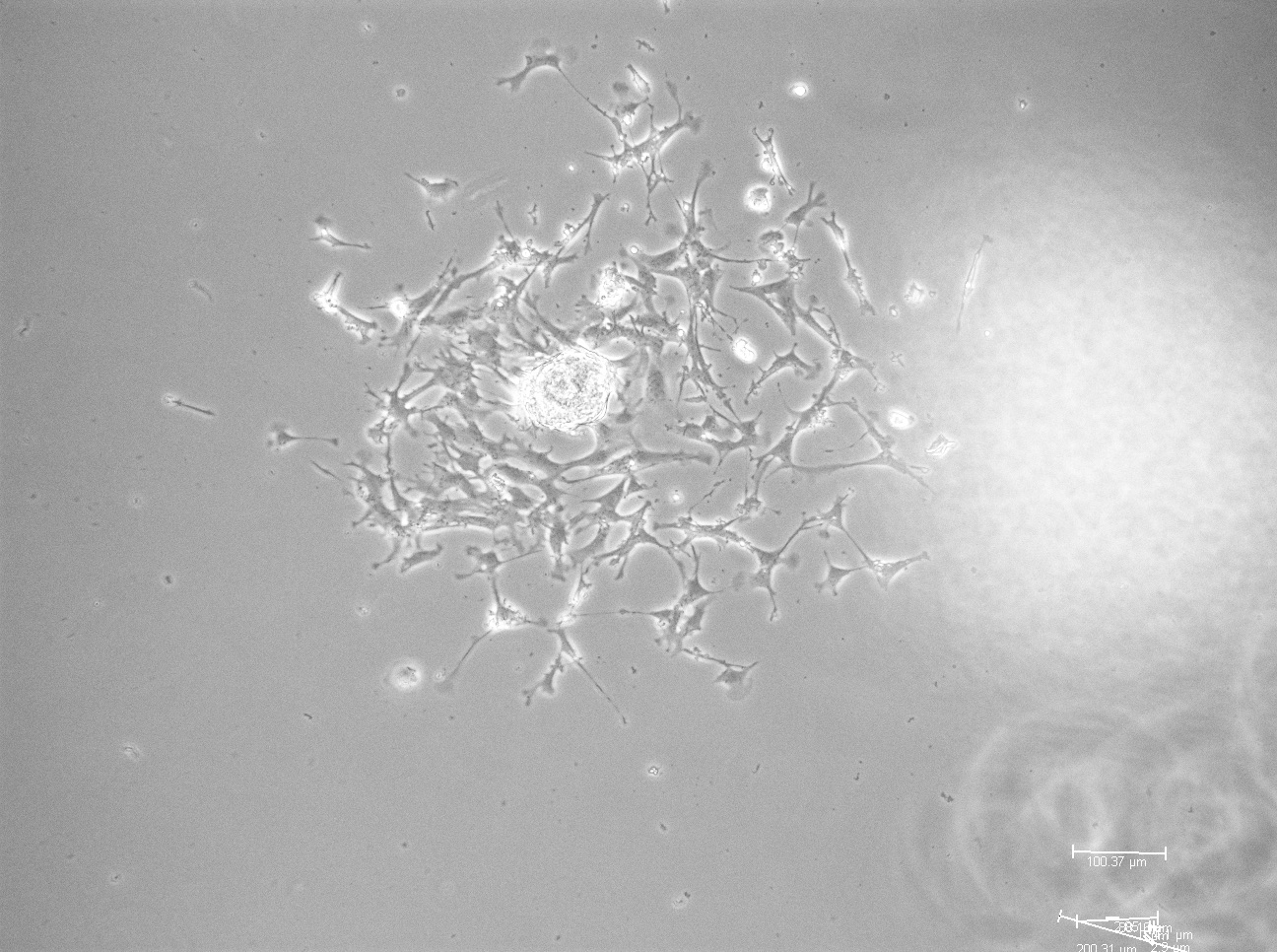 | 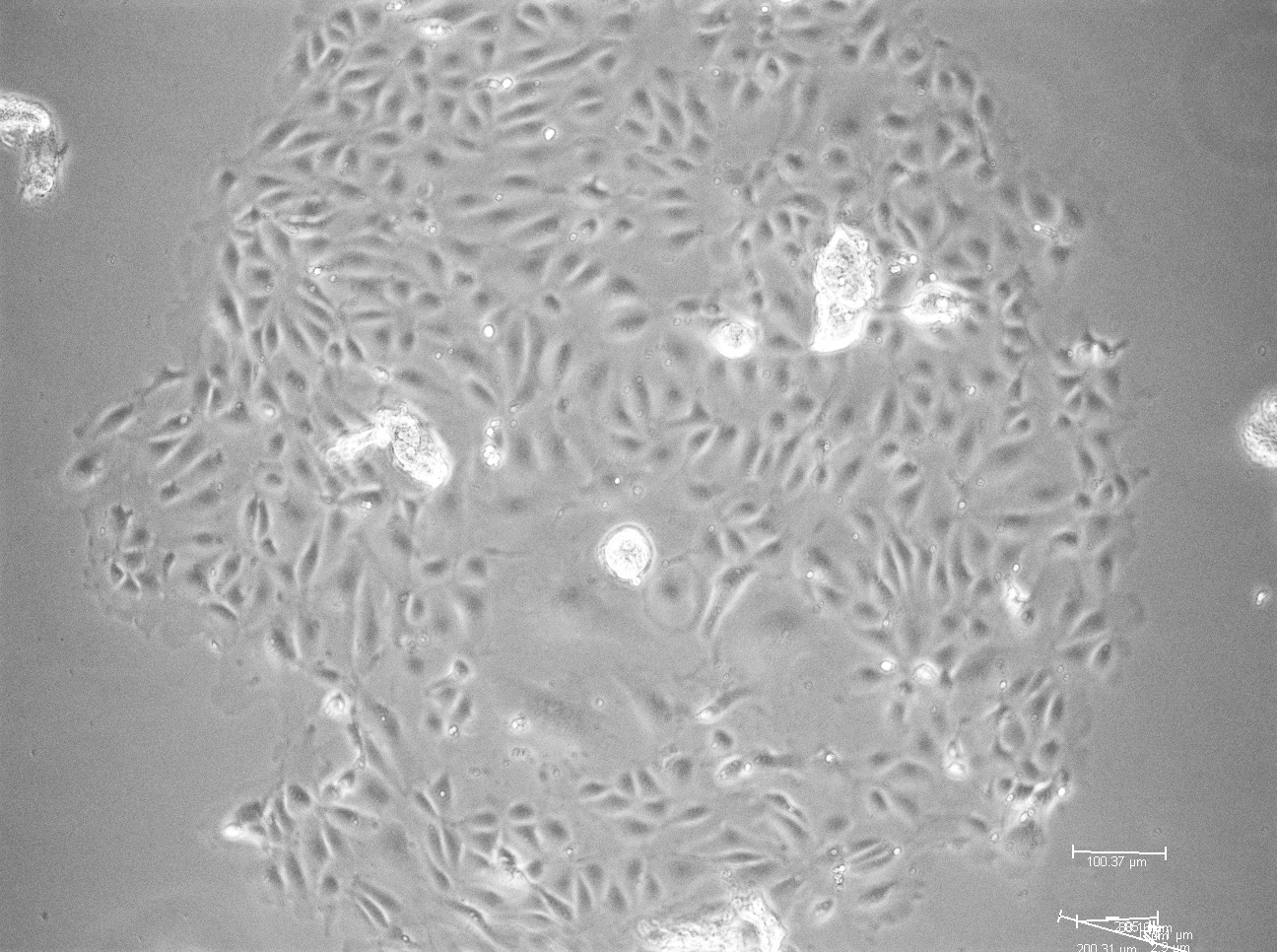 |
| --- | --- | --- | --- |
| 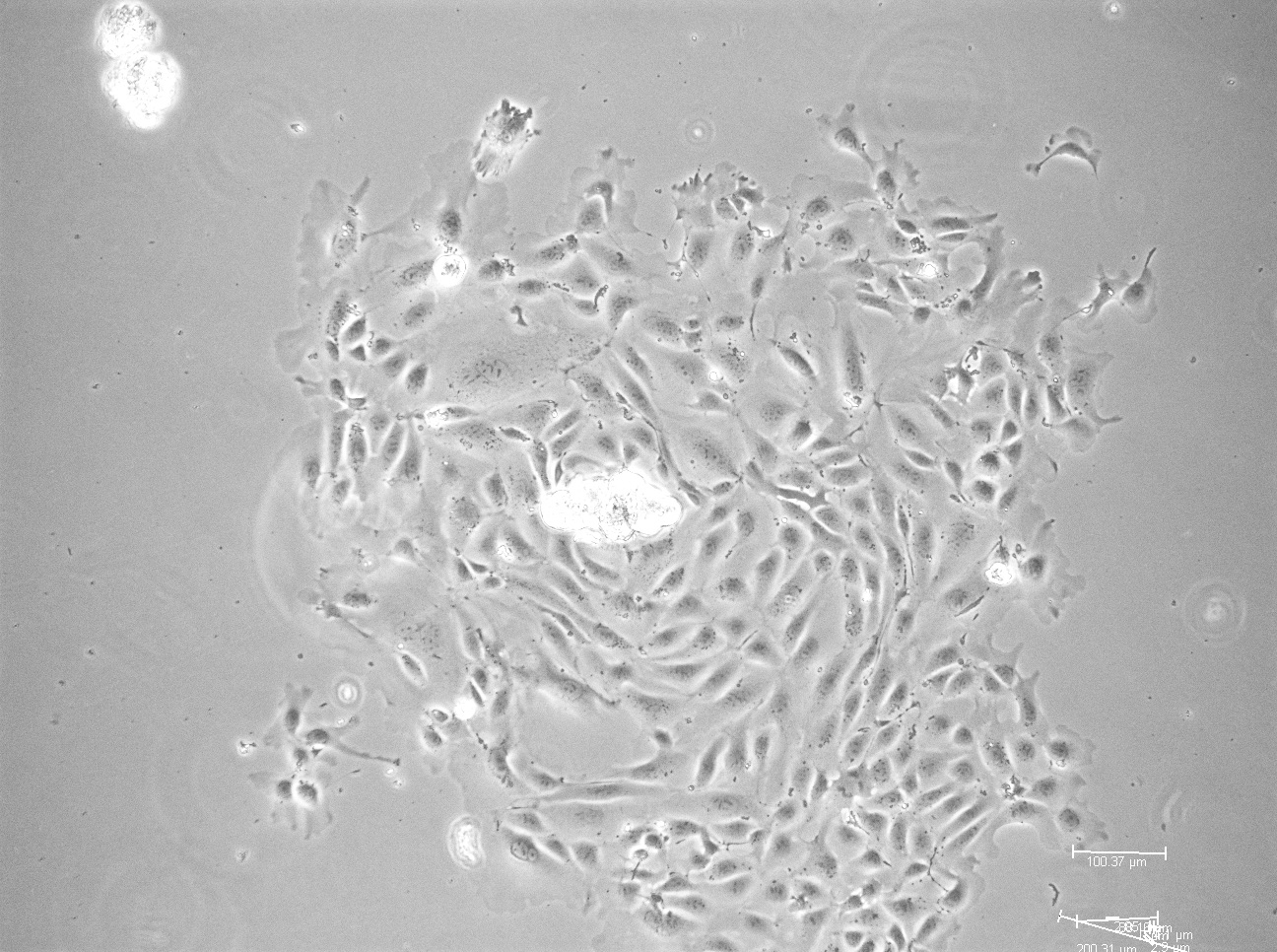 | 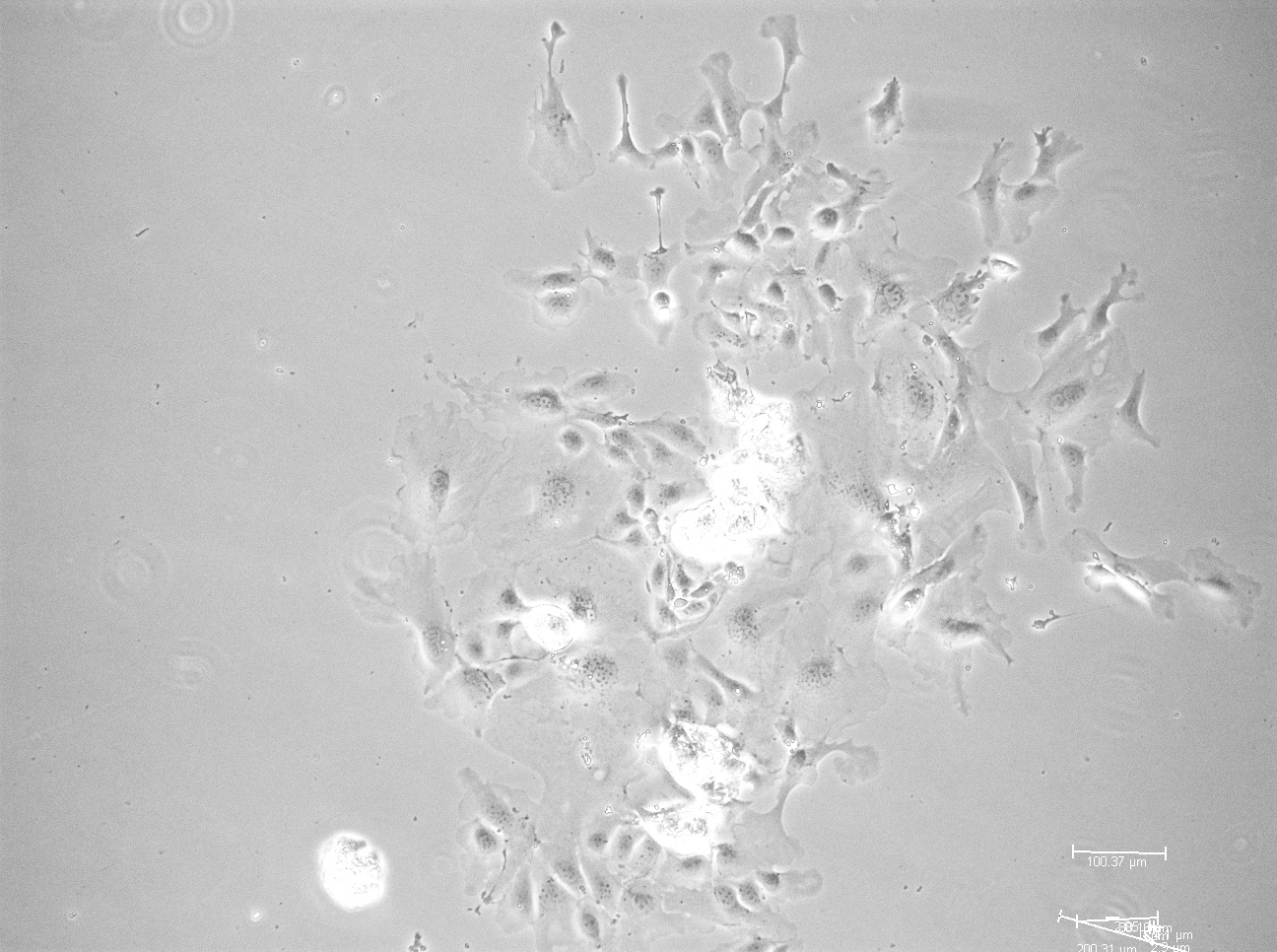 | 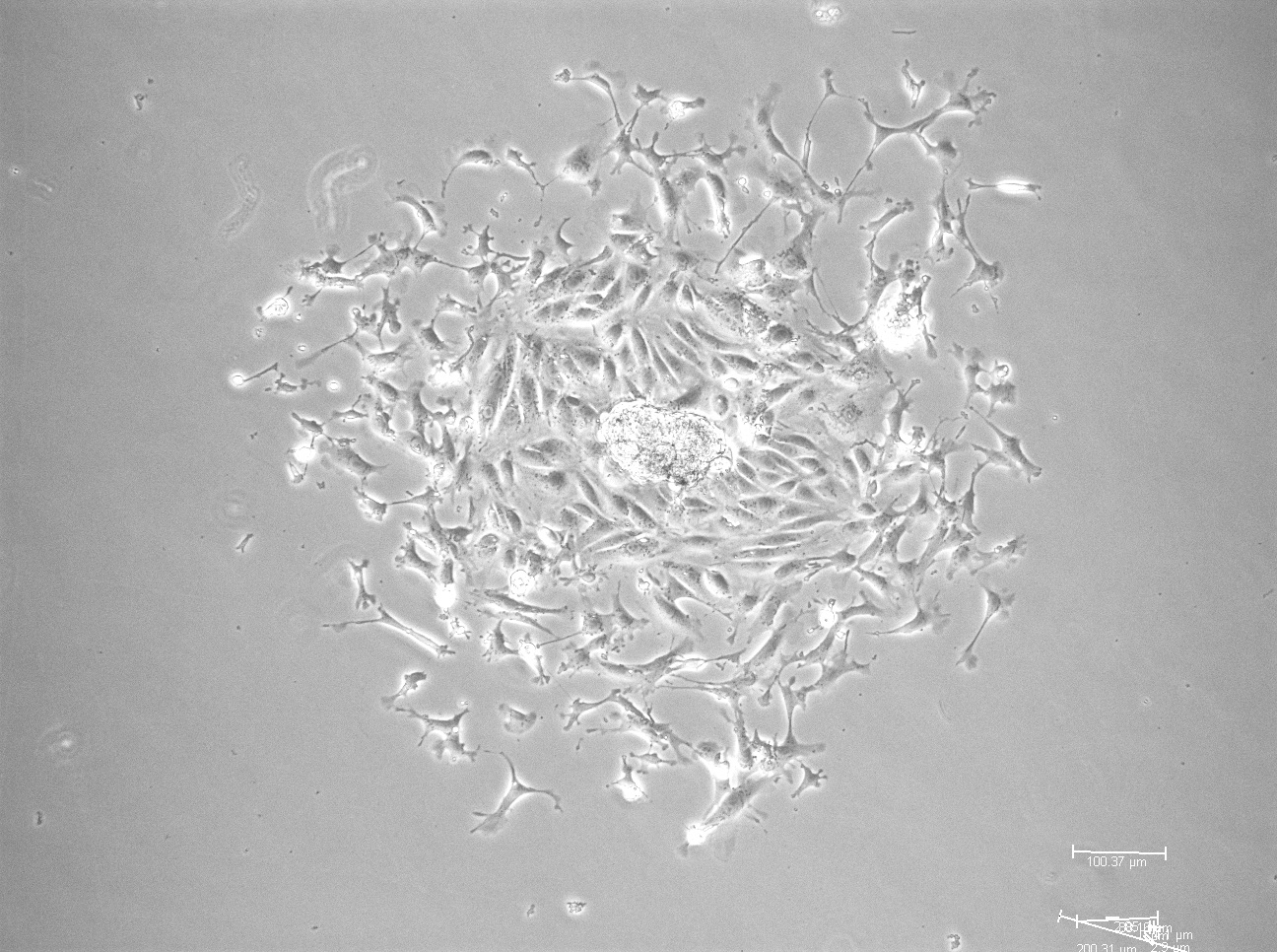 | 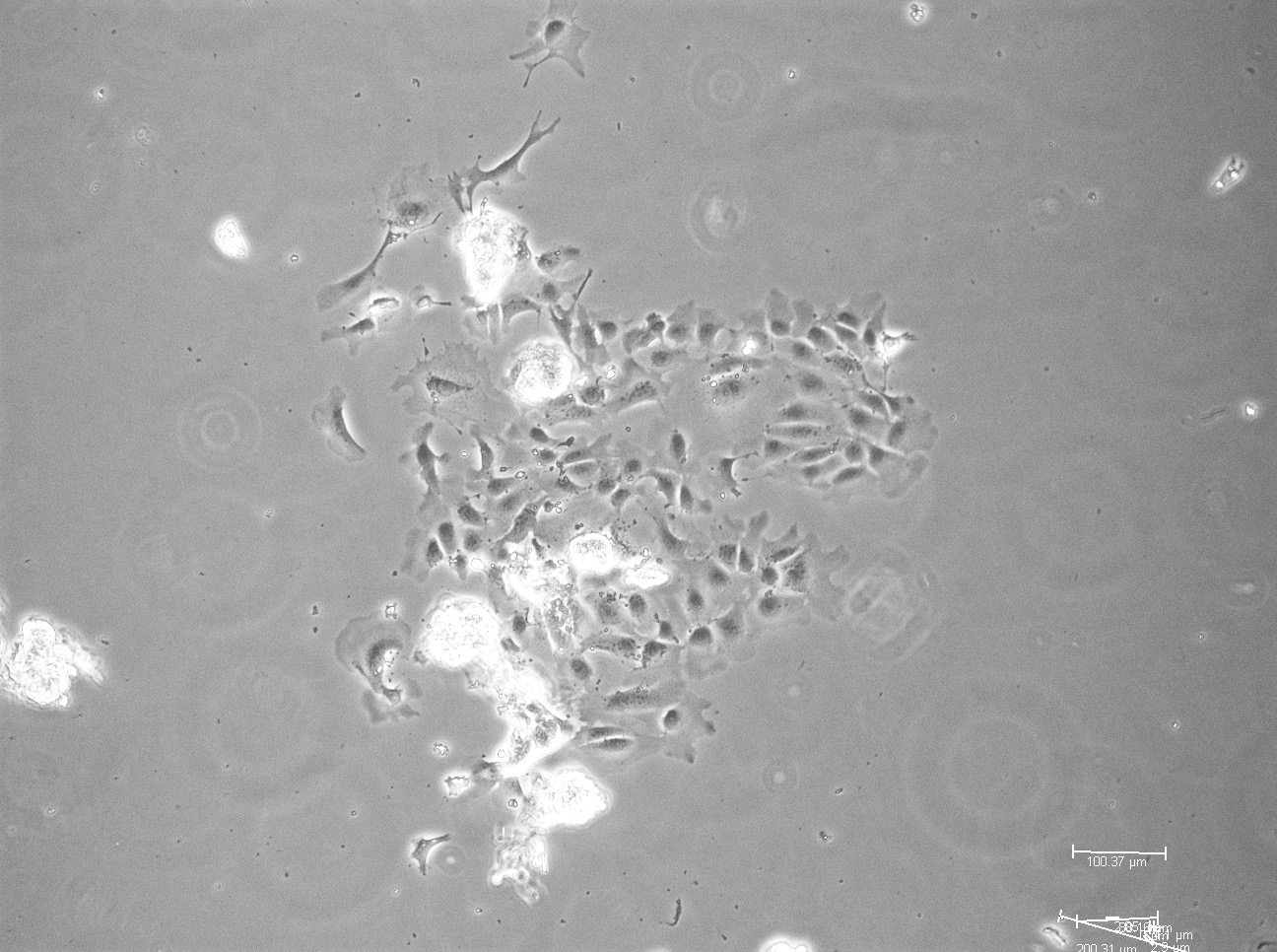 |
| 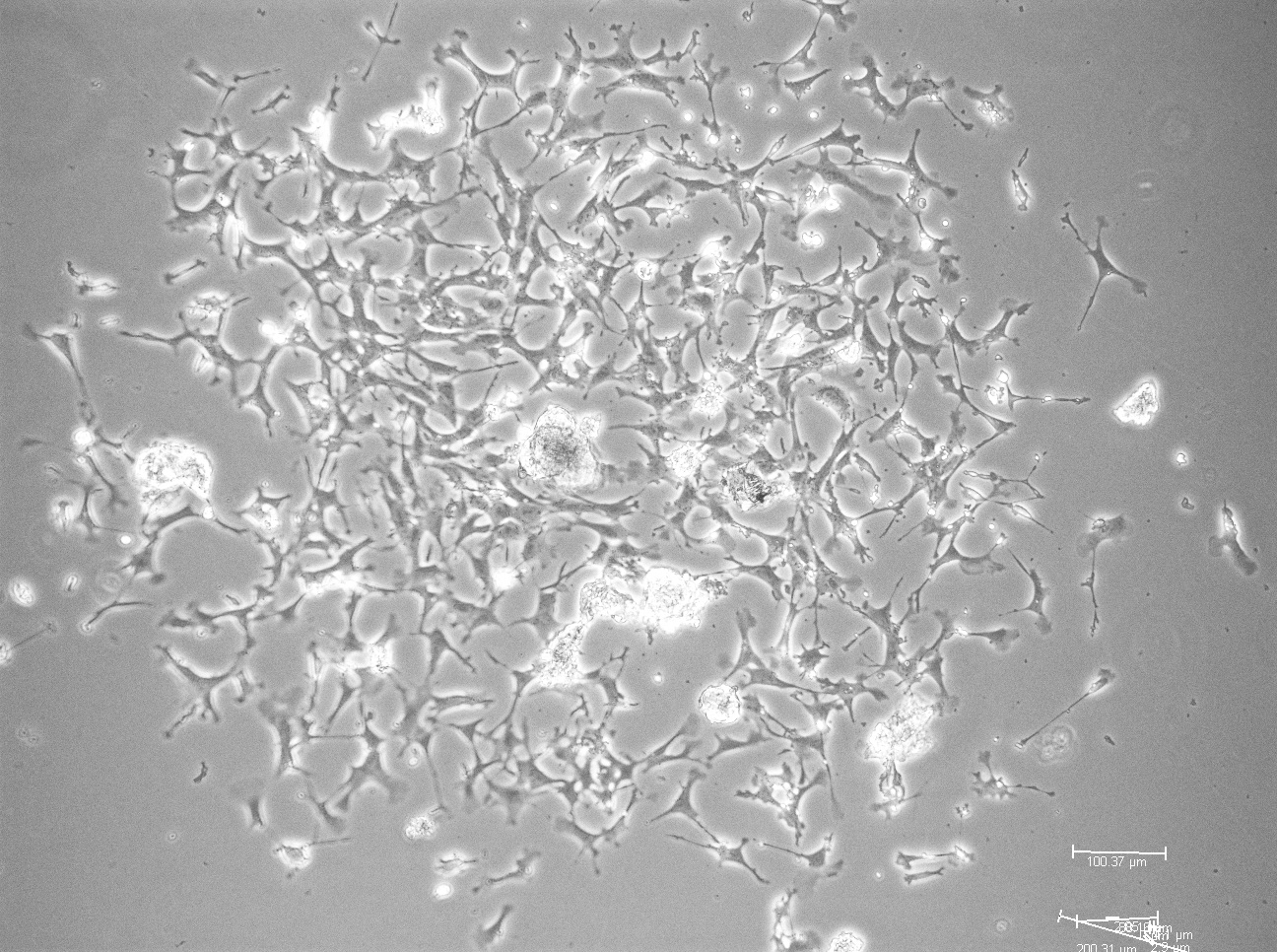 | 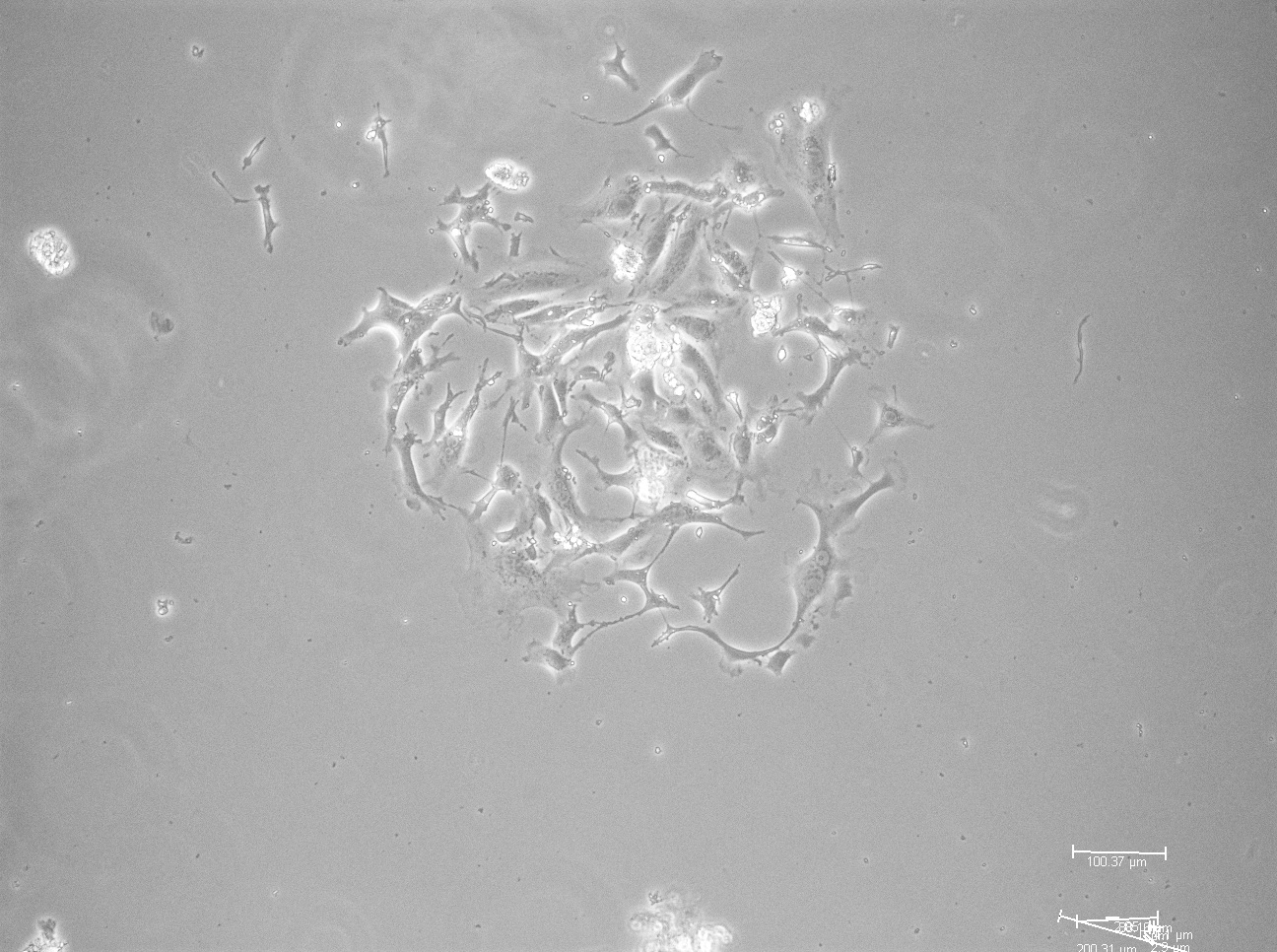 | 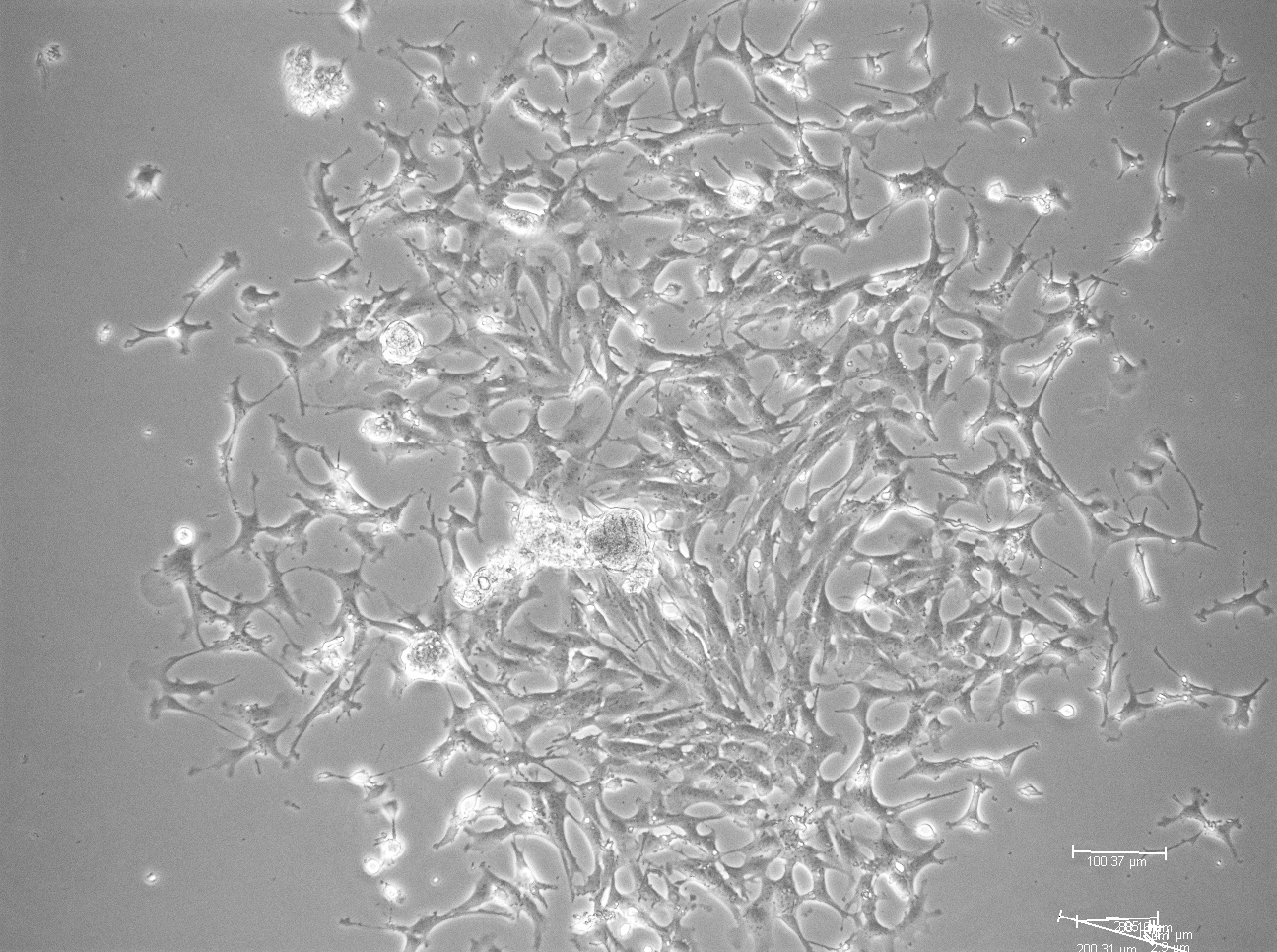 | 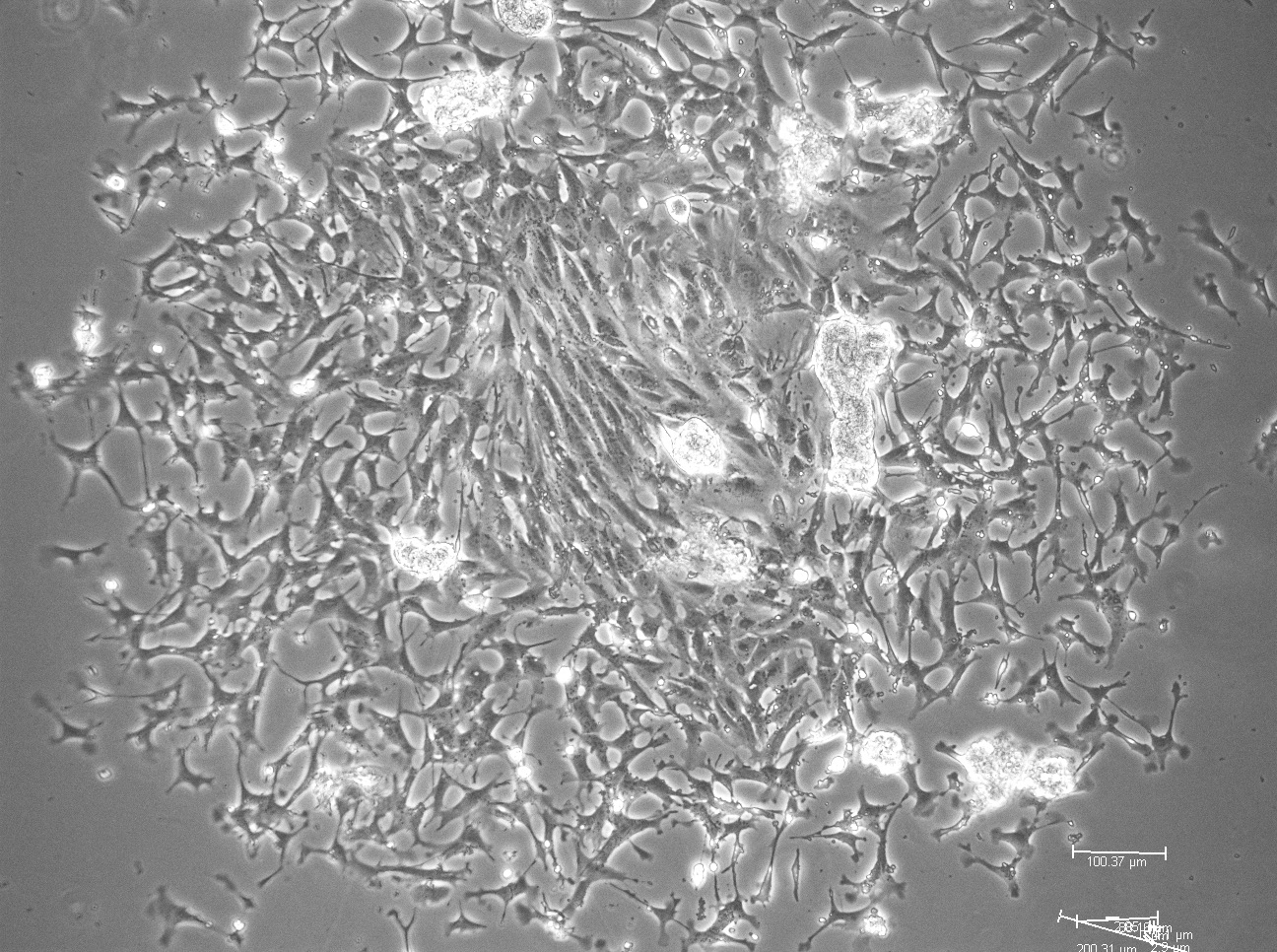 |

**B.**

| 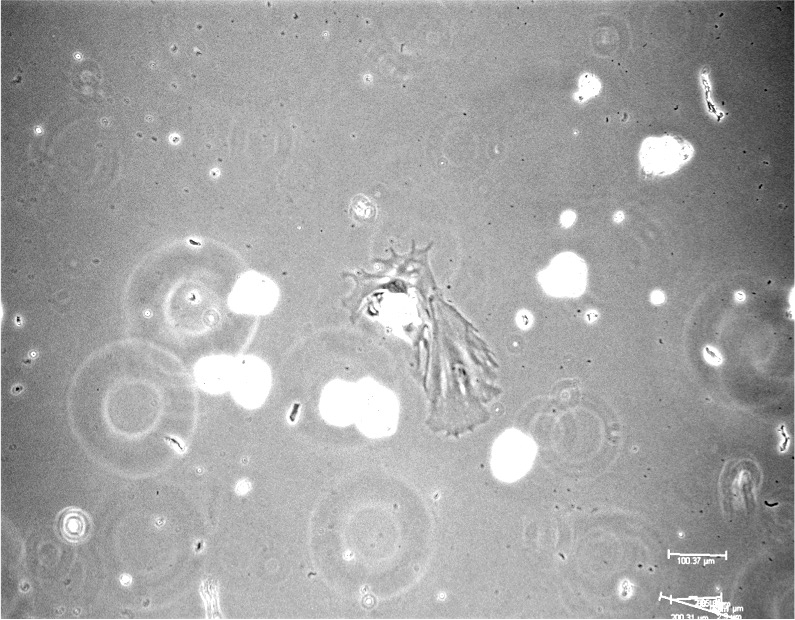 | 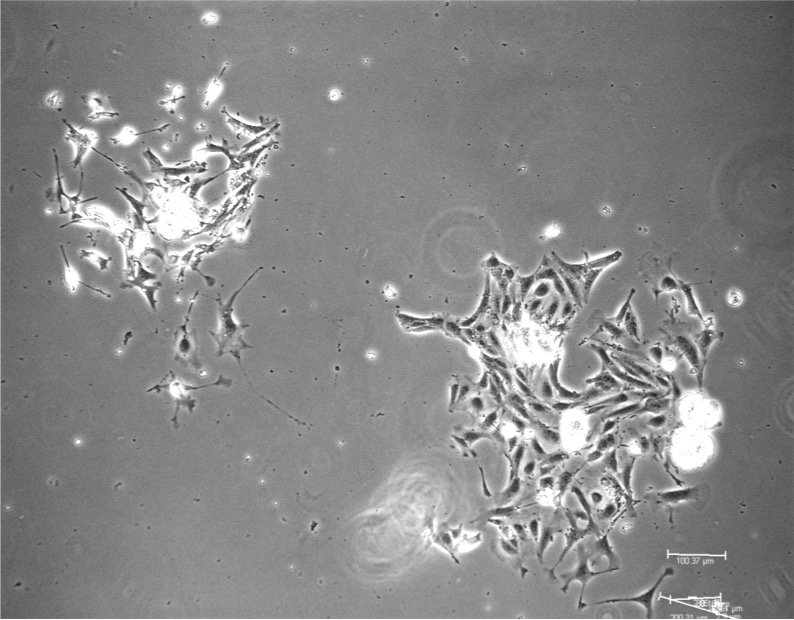 | 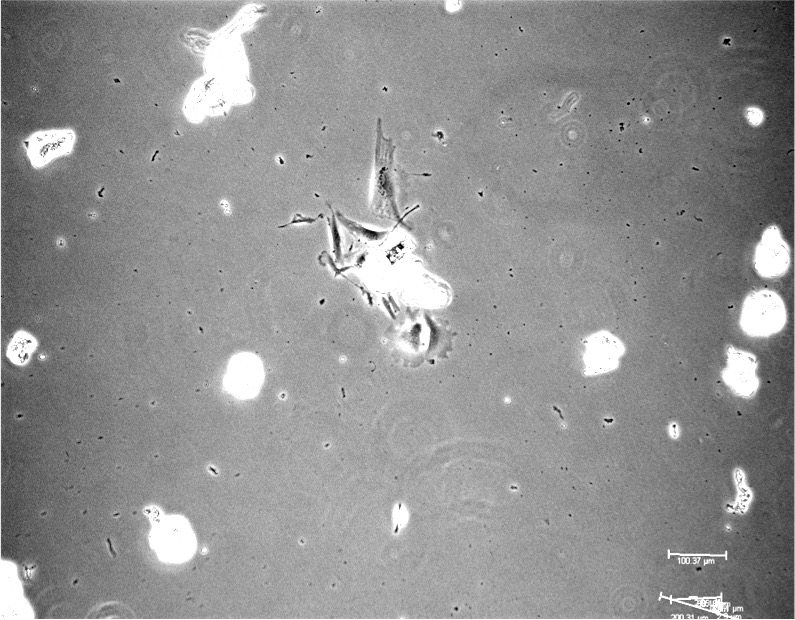 | 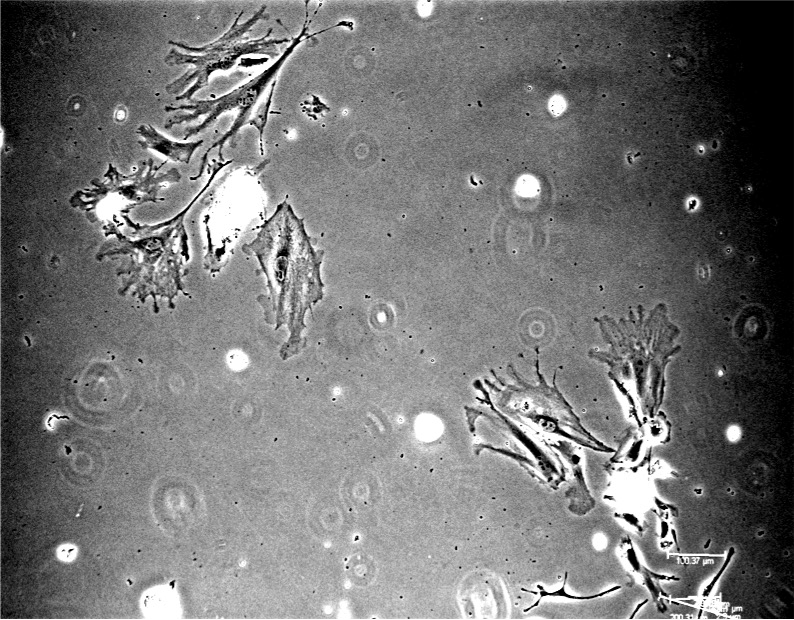 |
| --- | --- | --- | --- |
| 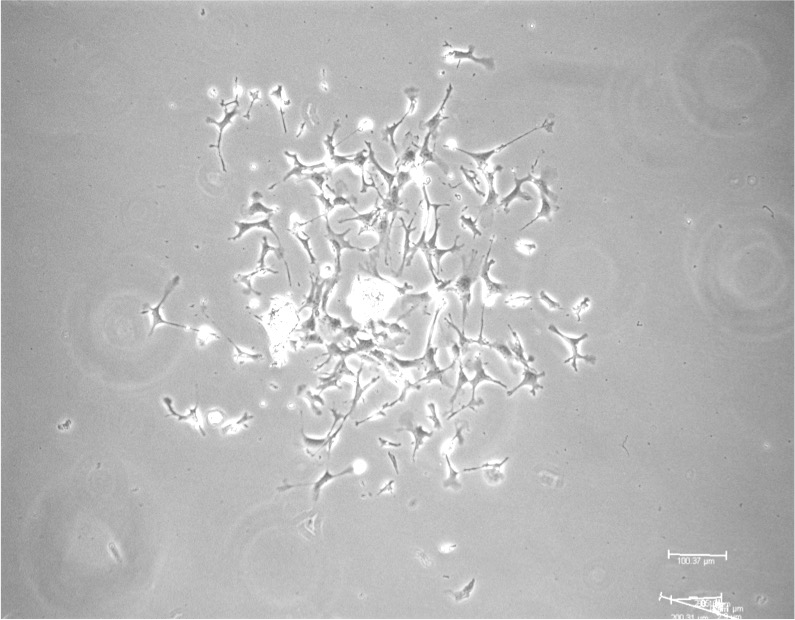 | 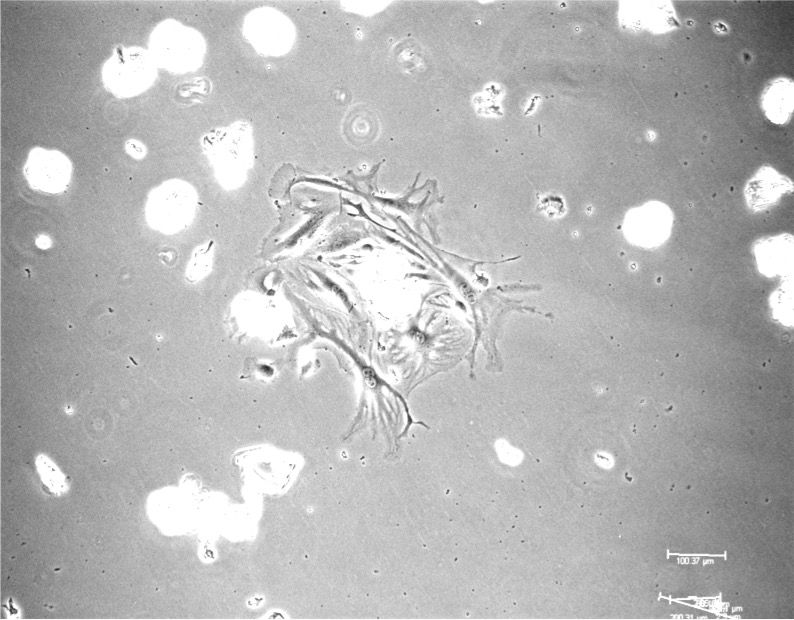 | 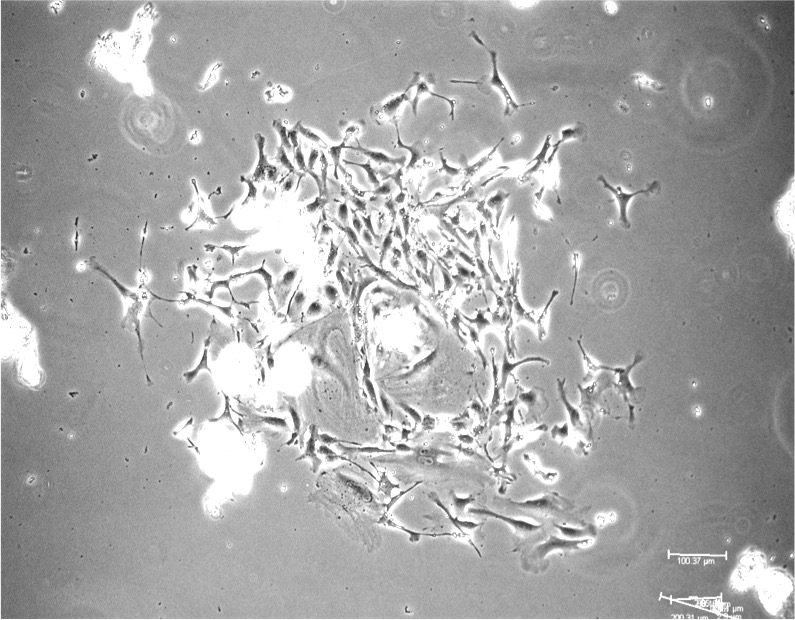 | 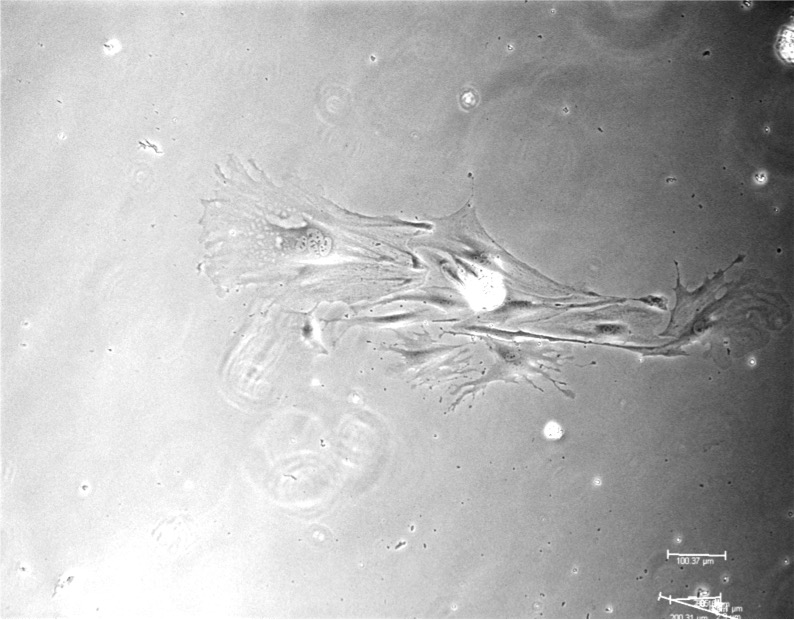 |
| 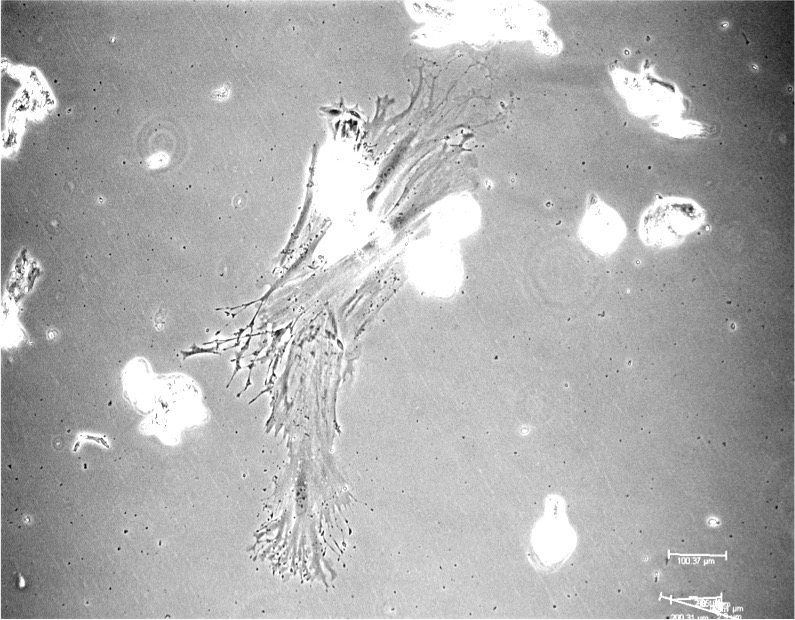 | 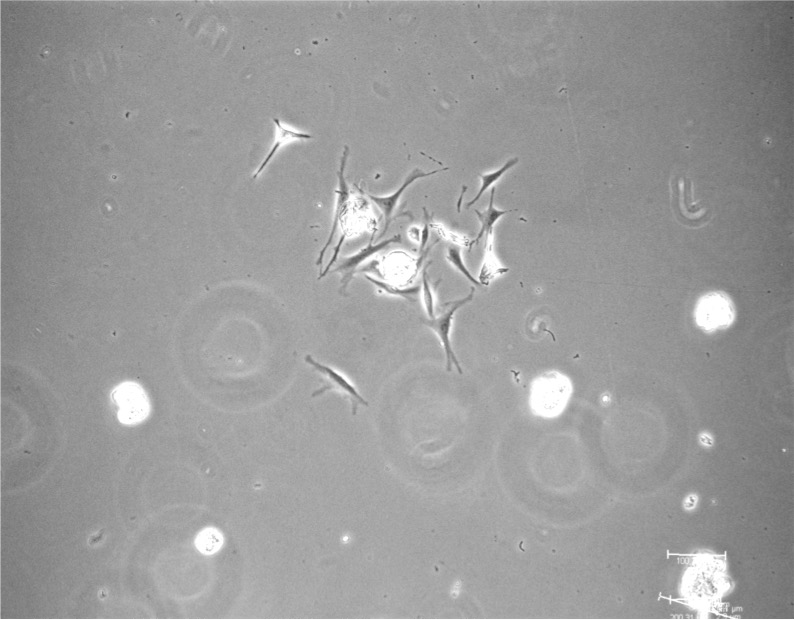 | 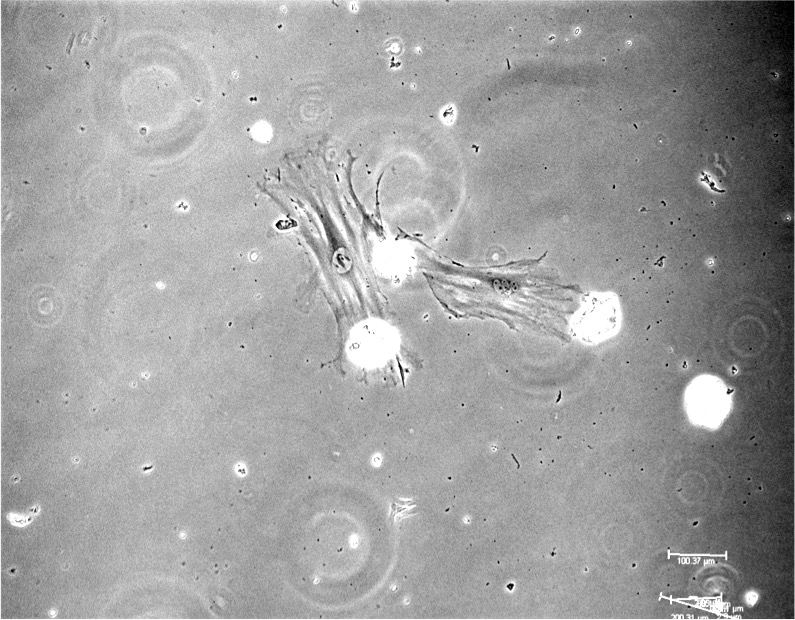 | 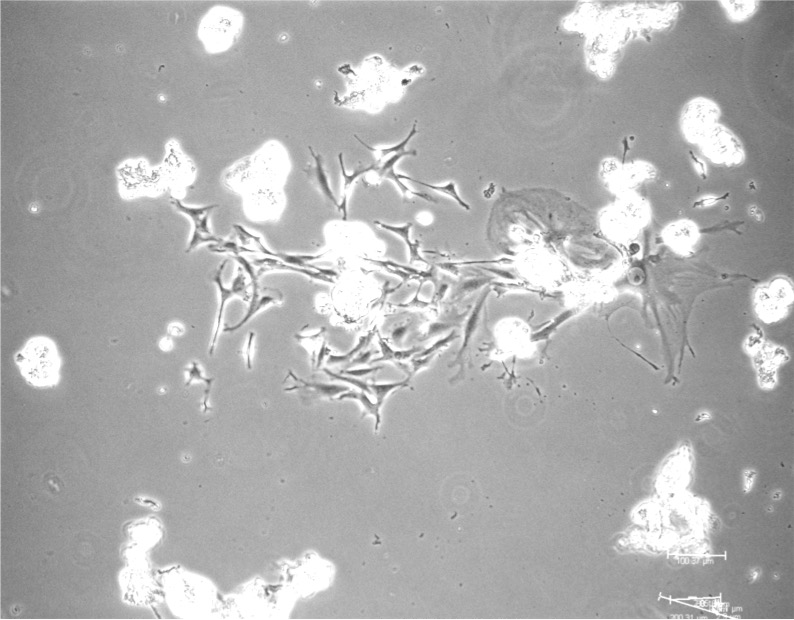 |

**Figure S4. Primary glomerular cell outgrowth colony at Day 5 in culture.** Live cell brightfield images of 12 separate patches of cells are shown. A) Every glomeruli shows cell outgrowth. WT glomerular cells that have moved out appear uniform in size. B) Numerous glomeruli from Alport KO mice have few outgrowths from glomeruli. Cells that have moved out from individual glomeruli are heterogeneous in size and morphology.

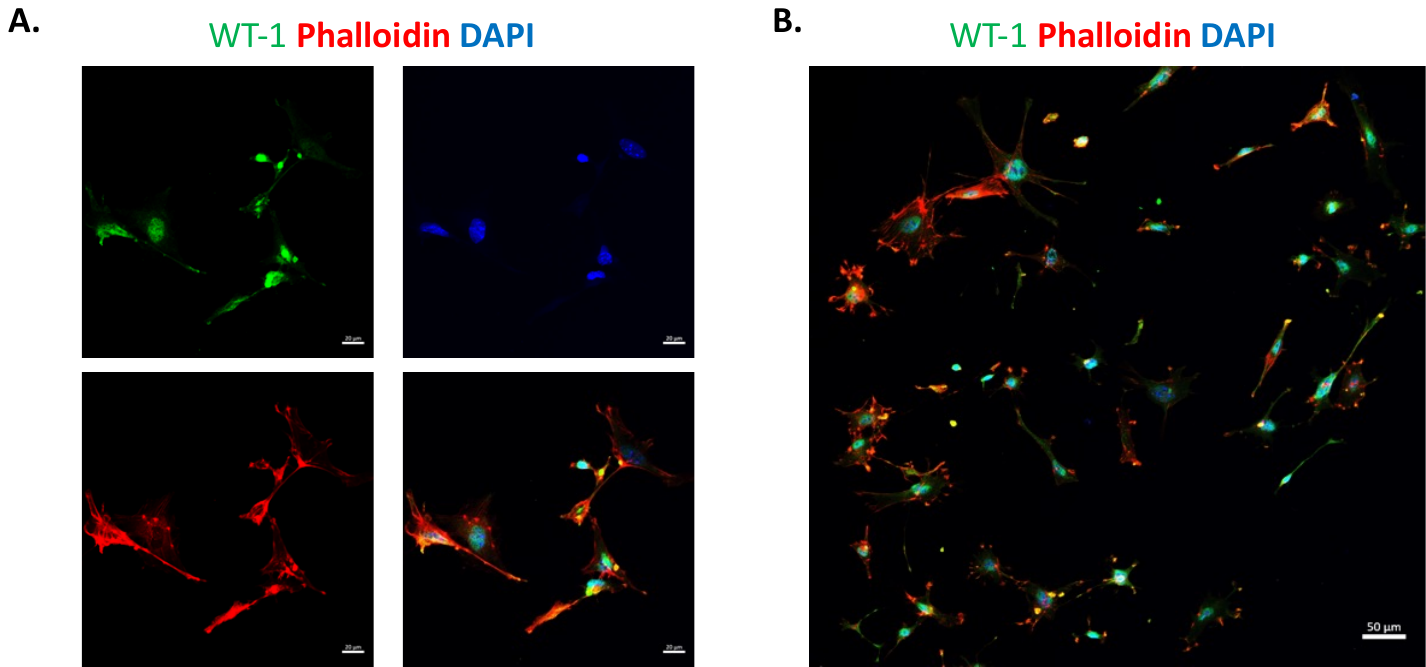

**Figure S5. Podocyte-specific WT-1 staining of reseeded glomerular outgrowth.** A. Podocytes stained for WT-1, Phalloidin, and DAPI are shown with merged image. B. Selected example of a low magnification image of primary podocytes identified by WT-1 positive staining, that was used to quantify that primary cells were >85% pure podocytes.

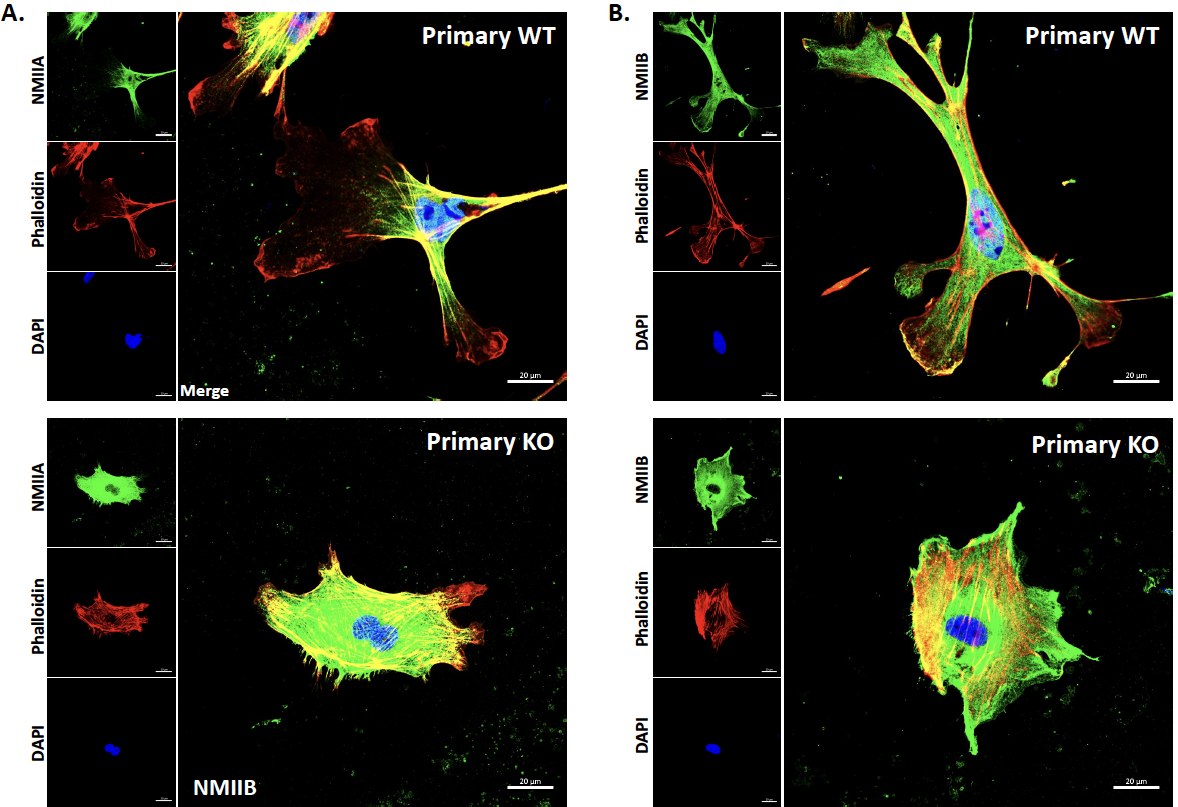

**Figure S6. Immunostaining of NMIIA/IIB in primary podocytes isolated from WT and Alport COL4α3 KO (KO) glomeruli.** Representative images of A) NMIIA and B) NMIIB co-stained with phalloidin to detect F-actin in primary podocytes from WT and KO mouse kidney. Left panels show the respective separated channel images for the enlarged merged image. NMIIA distribution is more centralized, and in both WT and KO cells, F-actin regions that are devoid of NMIIA are evident, whereas NMIIB distribution extends to F-actin positive edges of membrane extensions in both WT and KO podocytes. The scale bar represents 20 μm.

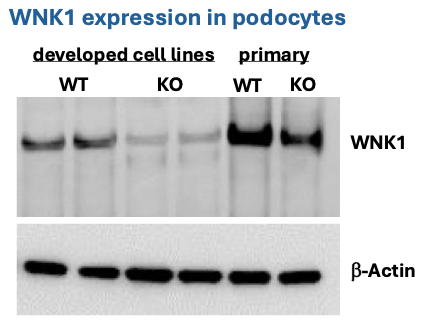

**Figure S7. Comparison of WNK1 expression in podocyte cell lines and mouse primary podocytes.**

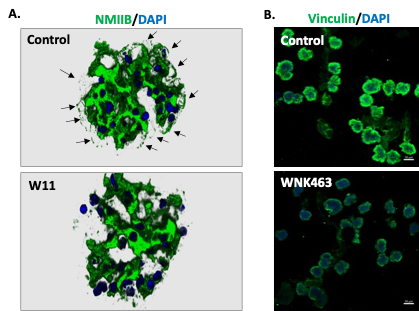

**Figure S8. Effects of WNK1 inhibition on NMIIB and vinculin distribution in isolated glomeruli. A) NMIIB localization shown in rendered 3D image of 9 slices (8um).** Arrows point to NMIIB-positive capillary loops where podocyte foot processes are located. B) Low magnification image of isolated glomeruli stained for vinculin after WNK1 inhibition *ex vivo*.

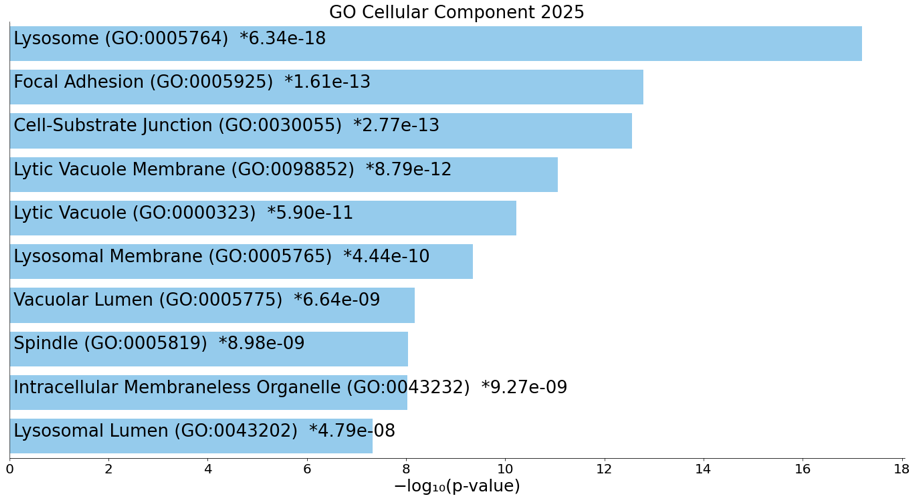

**Figure S9. Gene ontology analysis of differentially regulated genes in RNAseq comparison between WT and Alport KO primary podocytes, using Enrichr.** Total number of DEGs was 2758 after selection for genes with mean count >10, and -0.5>log2 fold change>0.5, N=1.
